# Stimulation-responsive enhancers regulate inflammatory gene activation through retention and modification of H2A.Z-variant accessible nucleosomes

**DOI:** 10.1101/2022.09.12.507565

**Authors:** Audrey Sporrij, Meera Prasad, Brejnev Muhire, Eva M. Fast, Margot E. Manning, Avik Choudhuri, Jodi D. Weiss, Michelle Koh, Song Yang, Robert E. Kingston, Michael Y. Tolstorukov, Hans Clevers, Leonard I. Zon

## Abstract

Prostaglandin E2 (PGE_2_) and 16,16-dimethyl-PGE_2_ (dmPGE_2_) are important regulators of hematopoietic stem and progenitor cell (HSPC) fate and offer potential to enhance stem cell therapies^1,2^. The mechanism of gene regulation in response to dmPGE_2_ is poorly understood. Here, we show that dmPGE_2_ regulates inflammatory gene induction by modulating the chromatin architecture and activity of enhancer elements in human HSPCs. We identified the specific genomic reorganization at stimuli-responsive enhancers that permits rapid transcriptional activation. We found that dmPGE_2_-inducible enhancers retain MNase-accessible, H2A.Z-variant nucleosomes that are permissive to binding of the transcription factor CREB. CREB binding to enhancer nucleosomes is concomitant with deposition of the histone acetyltransferases p300 and Tip60 on chromatin. Subsequent H2A.Z acetylation improves chromatin accessibility at stimuli-responsive enhancers. Our findings support a model where histone variant nucleosomes retained within inducible enhancers facilitate transcription factor (TF) binding. Acetylation of histone variant nucleosomes by TF-associated nucleosome remodelers creates the accessible nucleosome landscape required for immediate enhancer activation and gene induction. Our work provides a mechanism by which inflammatory mediators such as dmPGE_2_ lead to acute transcriptional changes and alter HSPC behavior.

## Introduction

Hematopoietic stem cells are characterized by their unique ability to self-renew and differentiate into all mature blood cell lineages. During normal homeostasis and in conditions of stress such as injury or inflammation, HSPCs maintain an appropriate balance of the hematopoietic system. HSPCs sense and respond to a variety of extrinsic signals that regulate their quiescence, proliferation, and differentiation^3,4^. A main mechanism of adaptation involves the activation of TFs that are downstream of signal transduction pathways to ensure appropriate transcriptional responses upon stimulation^5,6^. In higher eukaryotes, gene expression is regulated by the coordinated action of enhancers and promoters^7^. Stimuli-responsive TFs (STFs) tend to operate within the *cis*-regulatory repertoire that is established during cell fate specification and maintained by constitutive binding of lineage specific master TFs (MTFs)^8^. The access of STFs to these regulatory elements and their interaction with cofactors, such as transcriptional activators and chromatin remodeling complexes, depends largely on the local chromatin architecture^9^. Generally accepted features of active regulatory regions include open chromatin conformation, histone modifications and TF binding^10^. While promoters consist of a nucleosome depleted region that is established by chromatin remodelers, general TFs, and the basal transcription machinery, the typical chromatin organization and nucleosome configuration at enhancers remains unclear^11^.

Prostaglandins are physiologically active lipids produced in response to mechanical, chemical, or immunological stimuli. They sustain a variety of homeostatic and pathogenic functions. This includes roles in the inflammatory response. PGE_2_ is one of the most abundant prostaglandins produced in the body^12^. PGE_2_ and its stable derivative dmPGE_2_ act as important regulators of vertebrate HSPC development and homeostasis^1,13^. *Ex vivo* pulse exposure of HSPCs to dmPGE_2_ enhances engraftment and self-renewal in mice and clinical studies indicate benefits for hematopoietic stem cell transplantation (HSCT) outcomes in humans^2,14^. dmPGE_2_ predominantly exerts its effects by binding to the PGE_2_ receptor (EP) subtypes EP2 and EP4 on HSPCs^15^. Interaction with these G-coupled protein receptors enhances intracellular cAMP levels which activates signaling cascades and downstream effectors, for instance Wnt and β-Catenin^13^. Improved engraftment presumably results from upregulation of genes implicated in HSPC homing, such as *CXCR4*^16^. As enhancement of HSPC function by external stimuli supports a strategy to improve HSCTs, understanding the mechanism of gene regulation in response to inductive signals can provide a significant clinical opportunity.

Here, we sought to address how transcriptional induction is regulated during the HSPC response to dmPGE_2_. We exploited inducible TF binding to chromatin and identified, and then mechanistically dissected, enhancers controlling inflammatory gene expression changes in HSPCs. We assessed the chromatin accessibility and nucleosome organization of regulatory regions responsive to dmPGE_2_ and studied how the higher order chromatin structures changed following induction. We found that stimuli-induced enhancers retained MNase-accessible nucleosomes. These enhancer nucleosomes were enriched with the non-canonical histone variant H2A.Z and remodeled but not evicted during acute stimulation. Rather than prohibiting TF binding, we observed enrichment of the dmPGE_2_ responsive TF CREB at accessible nucleosomes within inducible enhancers. CREB binding is concomitant with deposition of the chromatin remodelers p300 and Tip60 that acetylates histone variant H2A.Z following dmPGE_2_ stimulation. This may further improve nucleosomes accessibility at stimuli-responsive enhancers and allow for binding of additional TFs and co-activator complexes. We suggest the nucleosome organization at enhancers is not exclusively repressive to gene regulation but favors STF binding, which enables rapid enhancer activation and inflammatory gene induction.

## Results

### CREB regulates gene expression changes through binding at enhancer elements

We sought to define the molecular mechanisms that underlie the transcriptional response of HSPCs to dmPGE_2_. We exposed human mobilized peripheral blood CD34^+^ HSPCs to 10μM dmPGE_2_ for 2 hours and performed extensive gene expression and chromatin profiling (Figure 1A). Using RNA-sequencing (RNA-Seq), we identified a total of 687 consistent differentially expressed genes (DEGs) after 2 hours of dmPGE_2_ treatment, when compared to vehicle (DMSO) treated control cells (Figure 1B). The effect of dmPGE_2_ on gene expression was overwhelmingly stimulatory. More specifically, 535 genes were ≥ 1.5-fold upregulated. This includes a significant number of genes involved in cell migration and cell cycle regulation (Supplemental Figure 1A, 1C). Among the upregulated set were genes representative of dmPGE_2_/cAMP/PKA signaling, including *PDE4B* and *PTGS2*^17^; several chemokines and cytokines, such as *CXCL2* and *CXCL8*^18^; and genes known to restrict HSPC proliferation and differentiation, for instance *NR4A1* and *JUNB*^19,20^(Figure 1C, Supplementary Table 1). We validated the gene expression signature observed by RNA-Seq through RT-qPCR in HSPCs for representative genes (Supplemental Figure 1D). 152 genes showed a ≤ 0.67-fold decrease in expression (Supplemental Figure 1B, Supplementary Table 1). Among the repressed genes, we found enrichment of genes that regulate cell division, such as *HOXB4* and *CCNF*^21^. To evaluate the functional impact of gene expression changes in human HSPCs, we assessed cell migration after dmPGE_2_ treatment *in vitro*. By transwell migration assay, which served as a proxy for the previously observed engraftment phenotype^1^, we observed greater migration of HSPCs after exposure to dmPGE_2_ (Supplemental Figure 1E). These data showed that enhanced engraftment *in vivo* after dmPGE_2_ stimulation is, in part, driven by transcriptional induction of migration genes.

**Figure 1.**
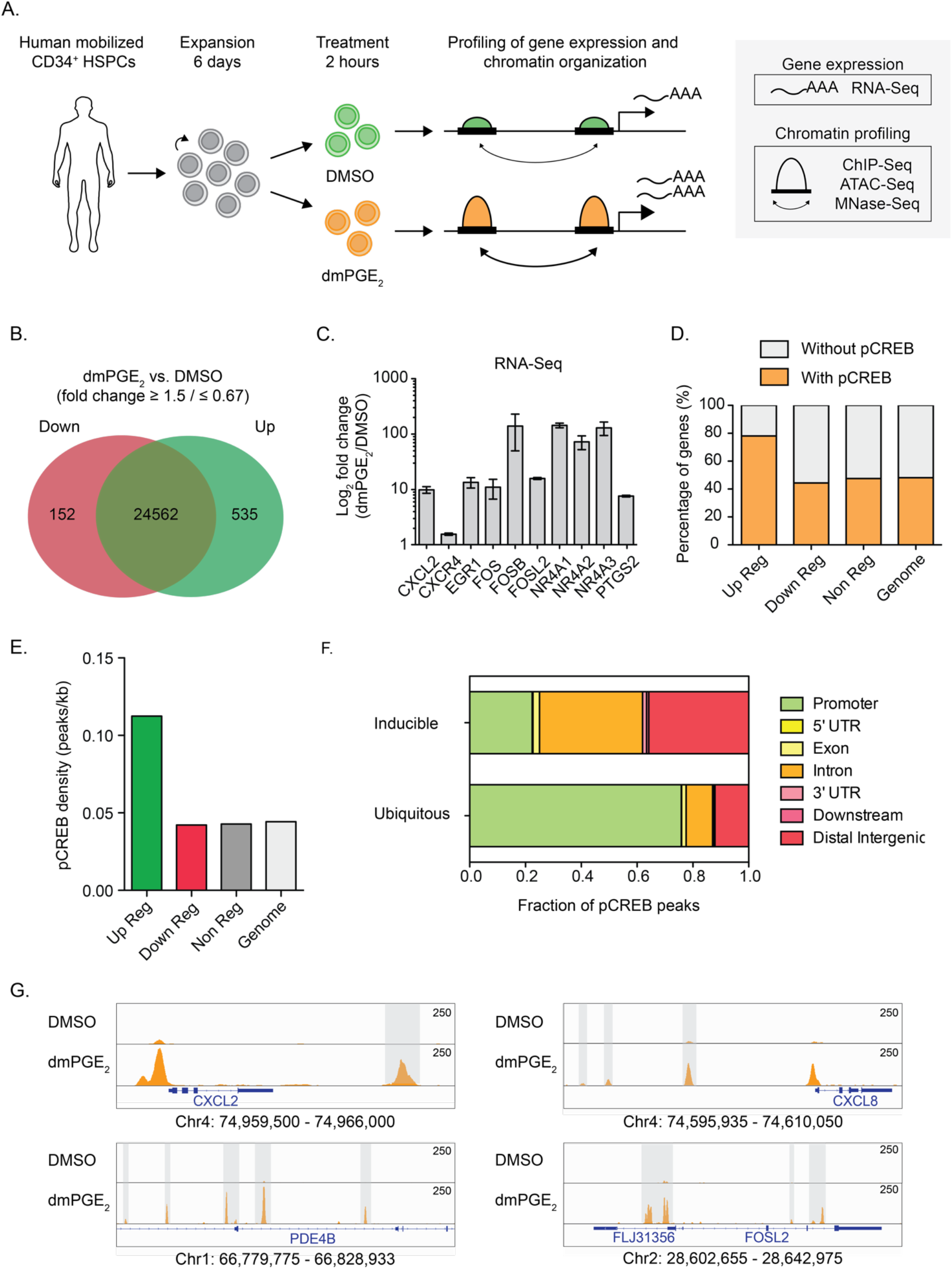
phospho-CREB regulates dmPGE_2_-induced gene expression changes through binding at distal regulatory elements. (A) Schematic representation of the experimental approach used in this study. Cells are stimulated for 2 hours with dmPGE_2_ or vehicle control (DMSO) after which transcriptome and epigenome profiling was performed. (B) Venn diagram showing the number of genes upregulated in green (535) and downregulated in red (152) in CD34^+^ HSPCs after 2 hours of dmPGE_2_ stimulation in comparison to control treated cells, as determined by RNA-Seq analysis. DEG criteria: FPKM ≥ 1 after treatment; fold change ≥ 1.5 or ≤ 0.67 (n = 3 biologically independent experiments). (C) Examples of genes identified as differentially expressed by RNA-Seq (n = 3 biologically independent experiments; mean values ± SEM). (D) Number of genes containing at least one pCREB peak in the proximity after dmPGE_2_ stimulation. pCREB peaks were assigned to a gene when located within a window from −5kb upstream of the transcription start site (TSS) to +5kb downstream of the TTS was considered (n = 2 biologically independent ChIP-Seq experiments). (E) Correlation between pCREB binding and gene expression in response to dmPGE_2_. pCREB density was calculated by dividing the total number of pCREB peaks associated to each gene category (up-, down, and nonregulated genes) by the total amount of base pairs that this category occupies in the genome. pCREB peaks were assigned to a gene when located from +5 kb upstream of the TSS to +5kb downstream of the TTS. Peak density in the genome was calculated by considering random distribution of pCREB sites in the whole genome. (F) Genomic distribution of unique pCREB peaks (inducible; present only after dmPGE_2_ stimulation) versus ubiquitous pCREB peaks. (G) Enrichment of pCREB binding at 4 representative dmPGE_2_ response genes: CXCL2 (promoter, intergenic), CXCL8 (promoter, intergenic), PDE4B (intronic), and FOSL2 (promoter, intronic). Gray bars indicate intronic and intergenic pCREB peaks. Genomic location of presented window is indicated at the bottom of the panels.

The TF CREB has previously been associated with dmPGE_2_ signaling^22^. Indeed, our transcriptomic analysis revealed that 30% (158/535) of upregulated genes are known targets of CREB (Supplemental Figure 2A). CREB binds to cyclic-AMP response elements (CREs) near its target genes where it, upon phosphorylation at serine 133 (S133) by protein kinases, promotes the recruitment of co-activator proteins^23^. This increases transcription of CREB-dependent genes^24^. One of protein kinases that phosphorylates CREB at S133 is protein kinase A (PKA). We assessed S133 phosphorylation of CREB in HSPCs after dmPGE_2_ treatment. Western blot analysis revealed increased abundance of S133 phosphorylated CREB (pCREB) while total TF protein levels remain unaltered (Supplemental Figure 2B). To correlate pCREB with gene induction, we performed chromatin immunoprecipitation sequencing (ChIP-Seq) with an antibody against pCREB. We found 31,198 binding sites (FDR < 0.1%) in CD34^+^ HSPCs treated for 2 hours with dmPGE_2_ compared to 8,332 sites in DMSO (Supplemental Figure 2C, Supplementary Table 2). Correlation of pCREB occupancy to DEGs showed enrichment of pCREB at genes induced by dmPGE_2_ (Figure 1D). In fact, 79% of the upregulated genes contained at least one pCREB peak in the proximity, that is a window from −5kb upstream of the transcription start site (TSS) to +5kb downstream of the TTS, after dmPGE_2_ stimulation compared to 47% prior to treatment (Figure 1D). This value increased to 85% by extending the window up to −100kb from the TSS to +25kb from TTS (Supplemental Figure 2C). Besides a higher percentage of upregulated genes containing pCREB, the TF density was also >2 times higher at upregulated genes compared to noninduced genes (Figure 1E, Supplemental Figure 1D). We observed no clear correlation between the magnitude of the transcriptional response and the density of pCREB (Supplemental Figure 1F). Genes downregulated by dmPGE_2_ showed no additional enrichment in pCREB compared to noninduced genes. This data demonstrated that dmPGE_2_ regulates pCREB activity and genomic binding near transcriptionally induced genes.

We next assessed the genomic location of pCREB bound regions. 23,386 (75%) sites displayed unique pCREB enrichment (‘inducible’ pCREB regions) in dmPGE_2_ treated HSPCs compared to control treated cells. The other 7,812 (25%) pCREB sites were present in both control and stimulated HSPCs (‘ubiquitous’ pCREB regions). The majority of inducible pCREB sites were located distal to the TSS of genes, with a strong representation in intronic and intergenic sequences (Figure 1F, 1G). Ubiquitous pCREB sites were predominantly enriched in promoter regions. This indicated that pCREB binding at putative distal regulatory elements plays a pivotal role in upregulating dmPGE_2_ response genes.

### Inducible enhancers gain master and signaling transcription factor binding

To understand the epigenetic impact of dmPGE_2_ on distal regulatory elements, we determined the effects of a 2-hour stimulation on the state of enhancers in HSPCs. H3K27ac distinguishes active enhancers from primed and poised enhancer elements that are marked by H3K4me1 alone or with H3K27me3, respectively^25^. We performed chromatin immunoprecipitation sequencing (ChIP-Seq) for these classifying histone modifications. We assessed H3K27ac enrichment in control and dmPGE_2_ stimulated HSPCs and identified a total of 25,998 active distal regulatory elements (Supplementary Table 3). Enhancers that regulate stimulus responsive gene programs can be distinguished from other active enhancers by their specific increase in H3K27ac upon receipt of the stimulus^26–28^. Comparison of H3K27ac enrichment between control and dmPGE_2_ treated HSPCs allowed us to identify putative distal enhancers involved in the response to dmPGE_2_. We identified a total of 954 (3.7%) enhancers that gained significant enrichment in H3K27ac following dmPGE_2_ treatment (Supplemental Figure 3A-B, and Methods). A subset of these stimuli-inducible enhancers (498/954, 52%) were only detected as active regulatory regions after dmPGE_2_ stimulation. While these ‘de novo’ enhancers were depleted of H3K27ac prior to stimulation, their H3K4me1^high^/H3K27me3^low^ state indicated that de novo enhancers reside in a primed state prior to activation (Figure 2A-B). The other fraction of dmPGE_2_-inducible enhancers (456/954, 48%) displayed significant enhancement in H3K27ac enrichment after dmPGE_2_ treatment (‘enhanced’, Figure 2A, Supplemental Figure 3C). Since the number of noninducible enhancers (25,044 regions, 96.3%) is much larger than the set of inducible enhancers, we randomly sampled a comparable number of noninducible enhancers (‘background’, 486) to ensure that observed differences are not due to variations in the size of defined enhancer categories. To assess epigenetic transitions at inducible enhancers, we profiled genome accessibility by assay for transposase-accessible chromatin sequencing (ATAC-Seq). We found a profound increase in DNA accessibility at inducible enhancers compared to background enhancers after dmPGE_2_ stimulation (Figure 2A-B), which is a sign of active chromatin reorganization^29^. In addition, the mean ATAC-Seq signal prior to activation of de novo enhancers suggested preexisting, yet minimal, accessibility before stimulation. This revealed that dmPGE_2_ stimulation results in rapid activation of stimuli-responsive enhancers that is concomitant with increased chromatin accessibility.

**Figure 2.**
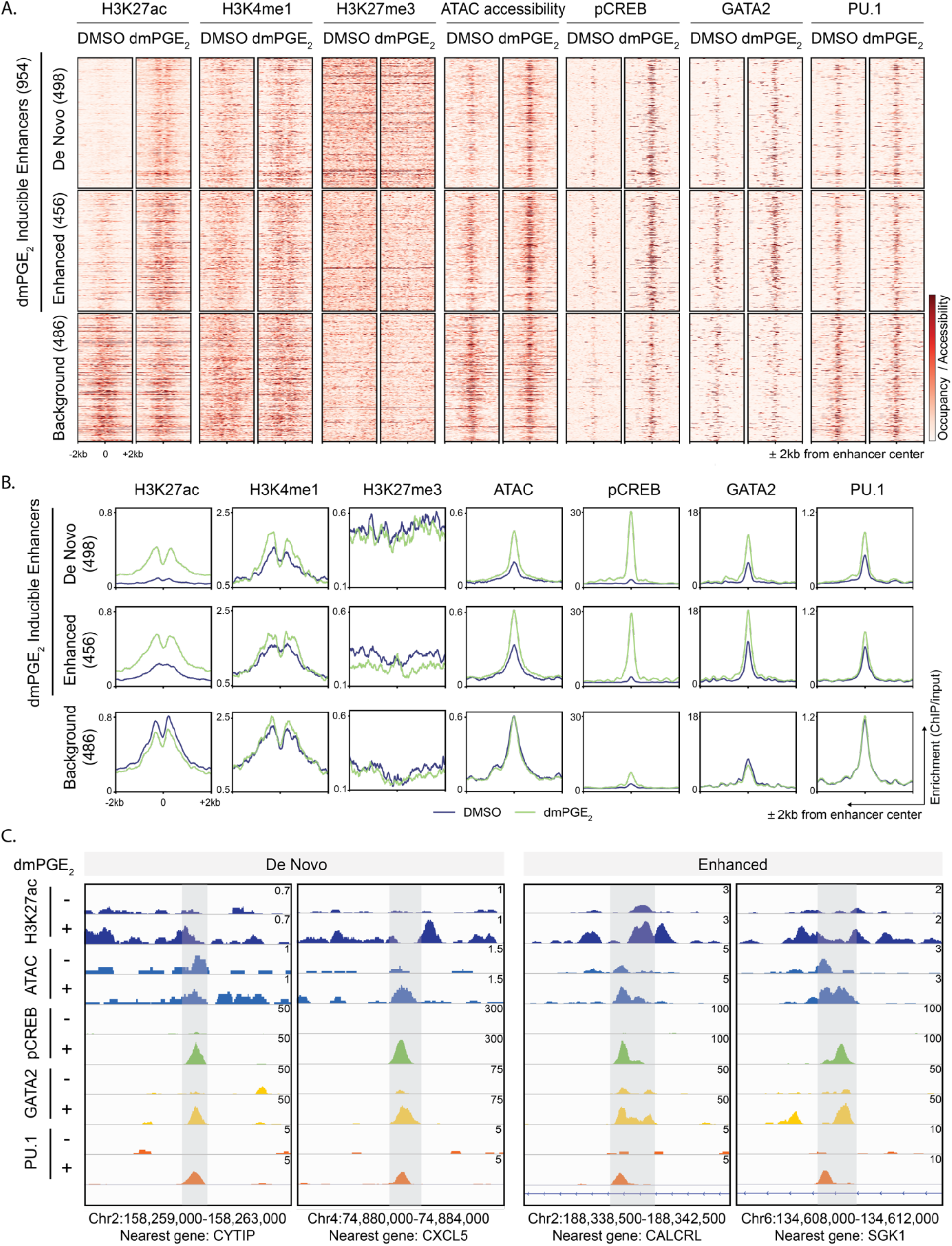
Stimuli-responsive enhancers gain chromatin accessibility and transcription factor binding after dmPGE_2_ stimulation. (A, B) Heat maps (A) and average enrichment profiles (B) of histone marks, ATAC accessibility and transcription factors at enhancers before and after dmPGE_2_ treatment. H3K27ac enriched regions identified by ChIP-Seq are classified as De Novo, Enhanced or Background enhancers according to the change in H3K27ac levels observed following dmPGE_2_ stimulation (n = 2 biologically independent experiments). A randomly sampled, comparable number of background enhancers (486) is shown. (C) Enrichment of histone mark, ATAC accessibility and transcription factor binding in response to dmPGE_2_ at 4 representative stimuli-response enhancers. Genomic location of presented window and nearest gene are indicated at the bottom of the panel.

It is well known that TFs act as anchors to recruit chromatin remodelers to regulate gene expression^7^. Most STFs do not possess pioneering activity and therefore preferentially bind DNA elements located within nucleosome depleted regions. Moreover, STFs often land at regulatory elements predefined by lineage specific MTF that have pioneer functions^27,30^. Given these insights, we assessed both pCREB occupancy and binding of the HSPC MTFs GATA2 and PU.1 at enhancers prior to and after dmPGE_2_ treatment. We found that pCREB colocalized with GATA2 and PU.1 at stimuli-responsive enhancers (Figure 2A, 2C). We moreover observed that inducible enhancers not only gain pCREB but also GATA2 and PU.1 enrichment after dmPGE_2_ stimulation (Figure 2B-C). No enrichment in MTF occupancy was observed in background enhancers (Figure 2A-B, Supplemental Figure 3E). These data showed that dmPGE_2_-inducible enhancers gain both STF and additional MTF deposition following stimulation.

The contribution of distal regulatory regions for transcriptional activation of dmPGE_2_ target genes is illustrated by the changes in the expression of genes regulated by stimuli-responsive enhancers compared to those regulated by background enhancers (Figure 3A, Supplemental Figure 4A). We tentatively assigned enhancers to individual nearest genes. To reasonably limit arbitrariness in gene assignment, we only considered genes with a mapped TSS within 15kb of an enhancer. The genes nearest to stimuli-inducible enhancers showed a greater transcriptional response than genes annotated to background enhancers (Figure 3A, Supplemental Figure 4A). This indicates that differential enhancer activity is directly reflected in gene expression levels. Importantly, genes associated with inducible enhancers belonged to several pathways, including cell signaling and blood cell migration such as *SGK1, CALCRL, CXCL2, CXCL5*, and *ITGA4* (Figure 2C, Supplementary Table 4). We found a clear correlation between the number of genes showing differential expression and the changes in enhancer activity. 16% of genes nearest to de novo enhancers display ≥ 1.5-fold induction after dmPGE_2_ treatment, compared to 9% of enhanced and 3% of background enhancers (Figure 3B, Supplemental Figure 4B). In total, 83 (16%) of the 535 upregulated genes were regulated by at least one stimuli-responsive enhancer (Figure 3C, Supplemental Figure 4C). This demonstrated that dmPGE_2_-mediated activation of stimuli-responsive enhancers correlates with modulation of gene expression.

**Figure 3.**
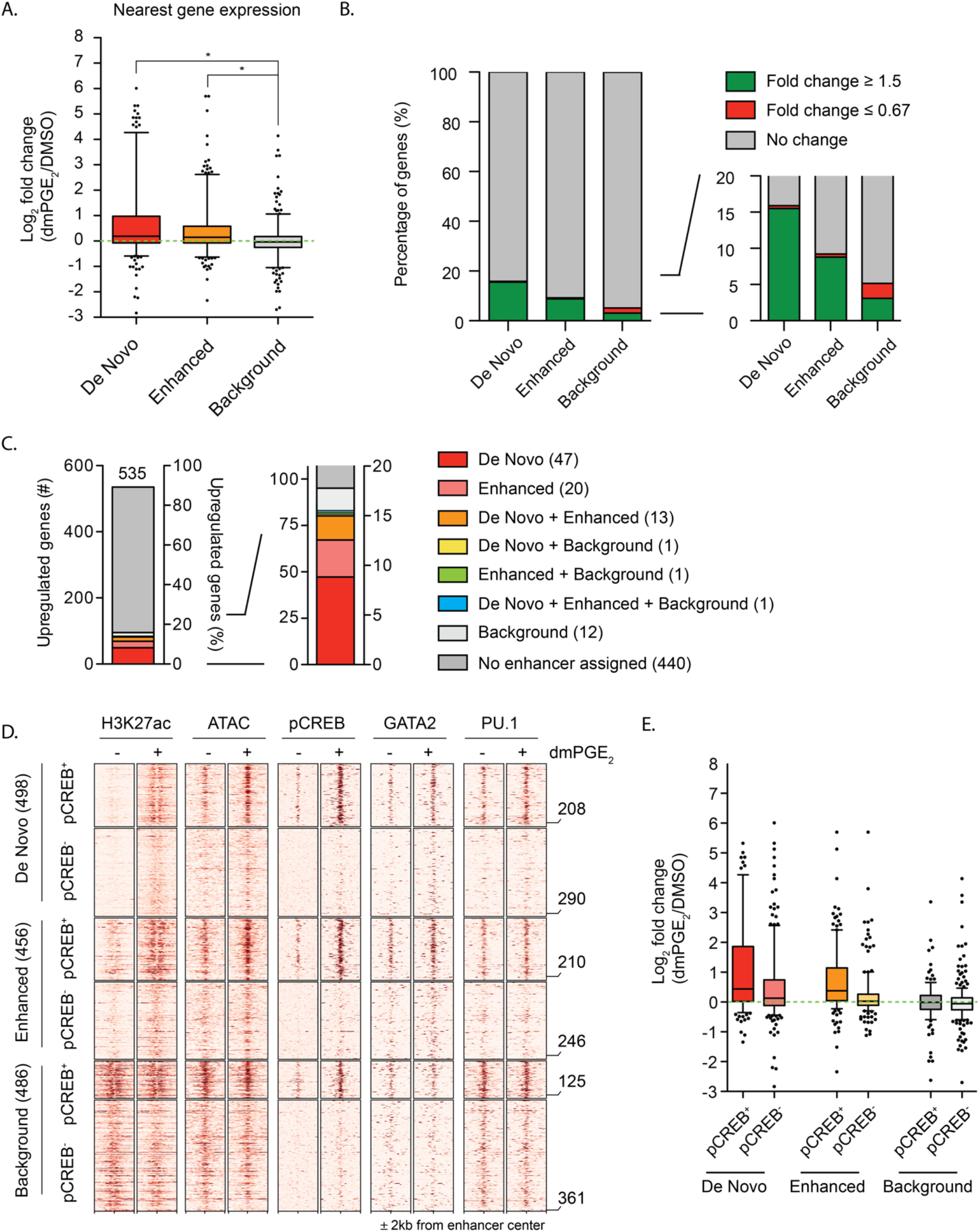
Stimuli-responsive enhancers mediate dmPGE_2_-induced gene expression changes. (A) Gene expression changes of genes associated with stimuli-responsive and background enhancers. Enhancers were assigned to an individual nearest gene. Only genes with a mapped TSS within 15kb of an enhancer were considered. Box plots shows median, 25^th^ and 75^th^ percentiles, whiskers are from 5^th^ and 95^th^ percentiles. Dots indicate outliers. (B) Percentages of enhancer nearest genes with fold changes in expression ≥ 1.5-fold or ≤ 0.67-fold for each enhancer category. (C) Upregulated genes with a fold change in expression ≥ 1.5 (535) and their associated enhancers. (D) Heat maps of H3K27ac, ATAC accessibility and transcription factors around enhancers before and after dmPGE_2_ treatment. De Novo, Enhanced or Background enhancers were subset based on the presence or absence of pCREB after dmPGE_2_. A randomly sampled, comparable number of background enhancers (486) is shown. Numbers of enhancers within each subset is indicated on the right of the heatmap. (E) Gene expression changes of genes associated with pCREB positive and pCREB negative enhancers. Enhancers were assigned to an individual nearest gene. Only genes with a mapped TSS within 15kb of an enhancer were considered. Box plots shows median, 25^th^ and 75^th^ percentiles, whiskers are from 10^th^ and 90^th^ percentiles. Dots indicate outliers. For all analysis presented here a randomly sampled set of background enhancers (486) was used.

To gain a better understanding of the role of pCREB at inducible enhancers, we further segmented enhancers based on the presence or absence of pCREB after dmPGE_2_ treatment. We found that 208/498 (42%) de novo, 210/456 (46%) enhanced and 125/486 (26%) background enhancers to contain at least 1 pCREB ChIP-Seq peak after dmPGE_2_ treatment. Chromatin accessibility and MTF binding increased more profoundly, but not exclusively, at pCREB^+^ stimuli-inducible enhancers (Figure 3D). The results support a prominent, but not restricted, role for pCREB in regulating gene expression through binding at inducible enhancers. Additionally, pCREB^+^ stimuli-responsive enhancer showed greater transcriptional changes than those without pCREB (Figure 3E, Supplemental Figure 4D). Together, our data suggested that increased enhancer chromatin accessibility and the cooperative binding MTFs and STFs at stimuli-responsive regulatory regions drives transcriptional induction.

### Inducible enhancers retain accessible nucleosomes after inflammatory stimulation

Various studies showed that TFs preferentially bind to sites of open, accessible chromatin^31^. However, recent work indicated that accessible chromatin may not necessarily represent nucleosome depleted regions^32,33^. To test whether greater chromatin accessibility at stimuli-responsive enhancers is due to nucleosome depletion, we performed micrococcal nuclease sequencing (MNase-Seq). This allowed us to map nucleosome positions and nucleosome occupancy changes after dmPGE_2_ stimulation. Because nucleosomes have varying sensitivities to enzymatic digestion, MNase titrations were performed to obtain a comprehensive map of the nucleosome landscape within a genome^32–34^. While comparing occupancy profiles between individual titration points provides insights on nucleosomal accessibility, the combinatorial analysis of all titration points within a given condition generates a complete view of the nucleosome organization^32^ (Supplemental Figure 5A). We prepared native nuclei from HSPCs treated with either DMSO and dmPGE_2_ and exposed them to increasing units of MNase. We selected 4 digestion points that generated increasingly larger fractions of mononucleosomal-size DNA. Mononucleosomal fractions comprised around 10%, 25%, 50%, and 75% of the input chromatin, respectively (Supplemental Figure 5B). Individual MNase titration profiles within a condition, as well as pooled average nucleosome occupancy profiles, revealed a TSS pattern similar to those previously reported^32^ (Supplemental Figure 5C-D). We observed lower average nucleosome occupancy at TSS-proximal regions of dmPGE_2_-responsive genes after treatment (Supplemental Figure 5E). This inverse correlation aligns with former studies which identified high levels of transcription to be concomitant with nucleosome eviction at the TSS, as elongation by RNA Polymerase II is thought to disrupt the nucleosomal organization^35,36^.

When assessing the nucleosome organization at inducible enhancers, we found that these regions presented in high nucleosome occupancy states when compared to active promoters (Figure 4A, Supplemental Figure 5E). Although high occupancy regions are traditionally thought of as ‘closed’, recent work indicated that accessible regulatory regions can retain nucleosomes^33,34^. We determined whether nucleosomes were repositioned or evicted at dmPGE_2_-responsive enhancers following stimulation. No decrease in average nucleosome occupancy was observed at stimuli-responsive enhancers. Inducible enhancers demonstrated higher nucleosome occupancy after dmPGE_2_ treatment (Figure 4A). This was not observed at background enhancers, indicating specificity of the phenomena to dmPGE_2_-inducible enhancers. Our data suggests that stimuli-responsive enhancers retained nucleosomes upon activation, rather than remodeled to a nucleosome free organization via nucleosome eviction.

**Figure 4.**
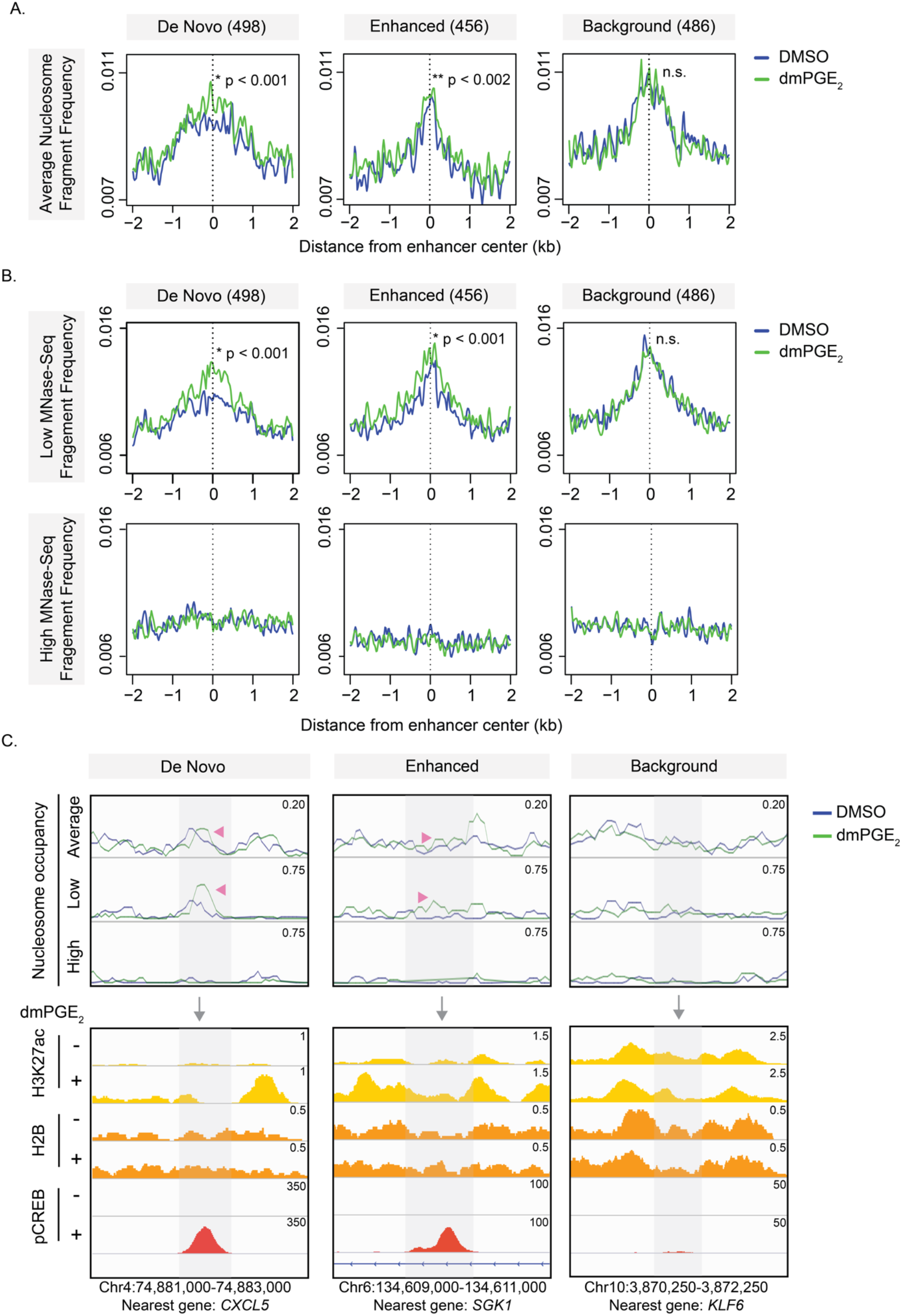
dmPGE_2_-responsive enhancers retain accessible nucleosomes after stimulation. (A) Average nucleosome occupancy profiles at stimuli-responsive and background enhancers from 4 MNase titration points. (n = 3 biologically independent MNase-Seq experiments). (B) Nucleosome profiles of low- and high MNase-Seq at stimuli-responsive and background enhancers (n = 3 biologically independent experiments). P-values by Wilcoxon rank-sum test. (C) Nucleosome fragment frequency and enrichment of H3K27ac, H2B, and pCREB at 3 representative stimuli-responsive and background enhancers. Genomic location of presented window and nearest gene are indicated at the bottom of the panel. Genomic location of presented window and enhancer nearest gene are indicated at the bottom of the panel. For all analysis presented here a randomly sampled set of background enhancers (486) was used.

Nucleosomes profiles from individual MNase titration points can be leveraged to determine how MNase sensitivity changes in specific regions (Supplemental Figure 5A). Light MNase conditions preferentially release accessible and unstable nucleosomes whereas stable nucleosomes are more resistant to enzyme digestion and only released at higher MNase concentrations^32–34^. We found greater low MNase-Seq signal at inducible enhancers after dmPGE_2_ stimulation (Figure 4B, upper panel) whereas high MNase digestion degraded the nucleosomal DNA fragments (Figure 4B, bottom panel). No changes in low MNase sensitivity were observed in background enhancers. Greater low MNase sensitivity after dmPGE_2_ treatment indicates a higher presence of MNase accessible nucleosomes. This shows that nucleosomes retained within inducible enhancers gained low MNase sensitivity of upon stimulation.

To ensure that the low MNase-Seq signal represents nucleosomes, we performed ChIP-Seq for the core histones H2B and H4 (Figure 4C, Supplemental Figure 6A). Inducible enhancers were not depleted of core histones prior to or after dmPGE_2_ stimulation. This is in line with our MNase-Seq data and suggested that nucleosomes are indeed present, and retained at, dmPGE_2_-responsive enhancers. In contrast to our expectations that increased chromatin accessibility at inducible enhancers (Figure 2B) resulted from nucleosome displacement or eviction, our MNase-Seq and ChIP-Seq data revealed that stimuli-responsive enhancers retained accessible nucleosomes upon activation.

Based on the accessible nucleosomes at enhancers, we next evaluated the relationships between nucleosomes and binding of pCREB at inducible enhancers. 10,169/23,386 (43%) of pCREB binding sites uniquely present after dmPGE_2_ treatment are located within the 25,998 active enhancers identified in HSPCs. We observed enrichment of phased nucleosomes at the summit of dmPGE_2_-unique pCREB peaks located within enhancers, both before and after dmPGE_2_ treatment (Figure 5A). To exclude the possibility that the MNase-Seq fragments observed at pCREB sites within enhancers represent non-histone proteins protecting from MNase digestion, we assessed fragment size distribution (Supplemental Figure 6B). We found a fragment periodicity that is characteristic for MNase-digested nucleosomes^37,38^. The majority of the MNase-Seq fragments that coincide with pCREB binding was 148 base pairs (bp) in length. This corresponds to precisely trimmed nucleosome core particles. Subnucleosomal peaks showed a clear ~10bp periodicity that reflects the accessibility of DNA as it is wound along the surface of the histone octamer^39^. This analysis suggested that MNase-Seq fragments mapping to pCREB sites within enhancers represent nucleosomes. Looking specifically at pCREB^+^ stimuli-responsive enhancers, we observed that the effects of dmPGE_2_ on nucleosome occupancy and low MNase sensitivity described earlier are further amplified at pCREB^+^ regions (Figure 5B-C). Assessment of individual loci confirmed pCREB binding at stimuli-responsive enhancers that show greater low MNase signal and increased nucleosome occupancies after dmPGE_2_ treatment (Figure 4C). Together, the data demonstrated that accessible nucleosomes are retained at dmPGE_2_-responsive enhancers and that these nucleosomes are not prohibitive of pCREB binding.

**Figure 5.**
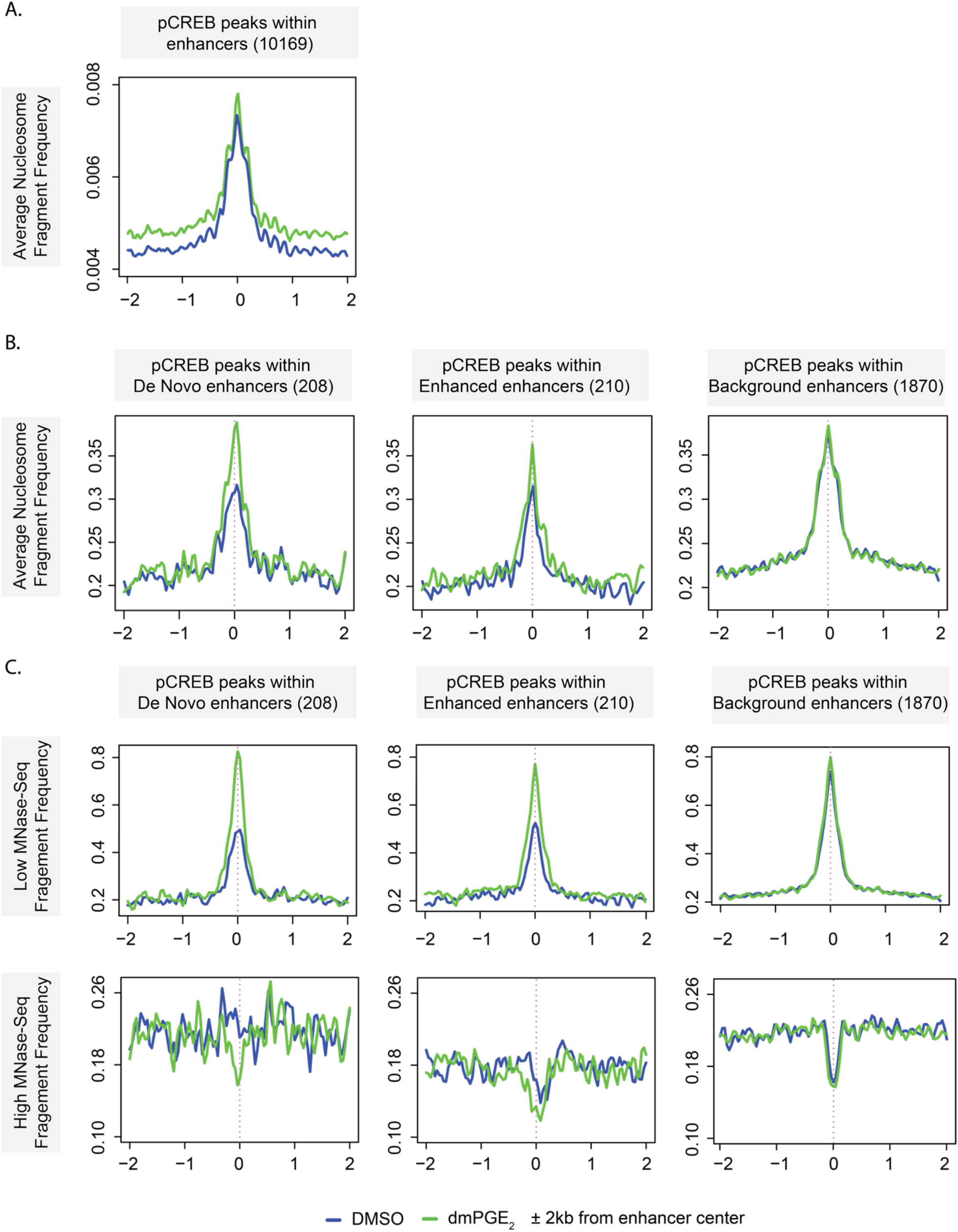
pCREB engages accessible nucleosomes in stimuli-responsive enhancers. (A) Average nucleosome frequency at pCREB peaks located within enhancers before and after dmPGE_2_ stimulation (n = 3 biologically independent MNase-Seq experiments). (B) Averaged nucleosome occupancy profiles at pCREB-bound sites within stimuli-responsive and background enhancers from 4 MNase titration points (n = 3 biologically independent experiments). (C) Nucleosome profiles of low- and high MNase-Seq at pCREB-bound sites within stimuli-responsive and background enhancers (n = 3 biologically independent experiments). For all analysis presented here a randomly sampled set of background enhancers (486) was used.

### Modification of H2A.Z-variant accessible nucleosomes at stimuli-responsive enhancers

We hypothesized that nucleosome at enhancers may exert important roles in pCREB binding. Nucleosome-driven TF binding has been observed for several other stress-responsive TFs^40^. To understand the conformational changes that underlie increased low MNase sensitivity of enhancer nucleosomes after dmPGE_2_ stimulation, we investigated other mechanisms that influence accessibility to nucleosomal DNA. Weakening of internucleosomal interactions by covalent modifications of histones as well as the introduction of histone variants increases DNA accessibility^41^. The histone variants H2A.Z and H3.3 can be incorporated in replication independent manners and are associated with enhancers^42–44^. Once incorporated, these non-canonical histones make for destabilized ‘fragile’ nucleosomes^45^. We assessed histone variant abundance at regulatory regions in the presence and absence of dmPGE_2_ stimulation through ChIP-Seq analysis for H2A.Z and H3.3. We observed a positive correlation between histone variant deposition and enhancer activity, with more profound presence of H2A.Z and H3.3 at higher H3K27ac levels (Supplemental Figure 7C-E). The histone variants were also found to occupy slightly different sites within enhancers. H2A.Z localized more central to enhancers the region where TFs bind, whereas H3.3-variant nucleosomes followed a more dispersed pattern and localized to the flanks of enhancers in a profile similar to H3K27ac (Supplemental Figure 7C-E). We noted minimal changes in H2A.Z enrichment following dmPGE_2_ stimulation. We did observe incorporation of the histone variant H3.3 in the nucleosomes of enhancers (Supplemental Figure 7C-E). When specifically assessing histone variants at pCREB^+^ stimuli-responsive enhancers, we found that pCREB binding directly overlaps with H2A.Z-variant nucleosomes (Figure 6A, Supplemental Figure 7A). To confirm interaction of pCREB with H2A.Z-variant nucleosomes, we performed complex immunoprecipitation (Co-IP) for pCREB and found association between the TF and H2A.Z histones (Figure 6B, Supplemental Figure 7B). These results indicated that pCREB overlaps with H2A.Z and suggest that DNA binding of the TF is not prohibited by H2A.Z-variant nucleosomes.

**Figure 6.**
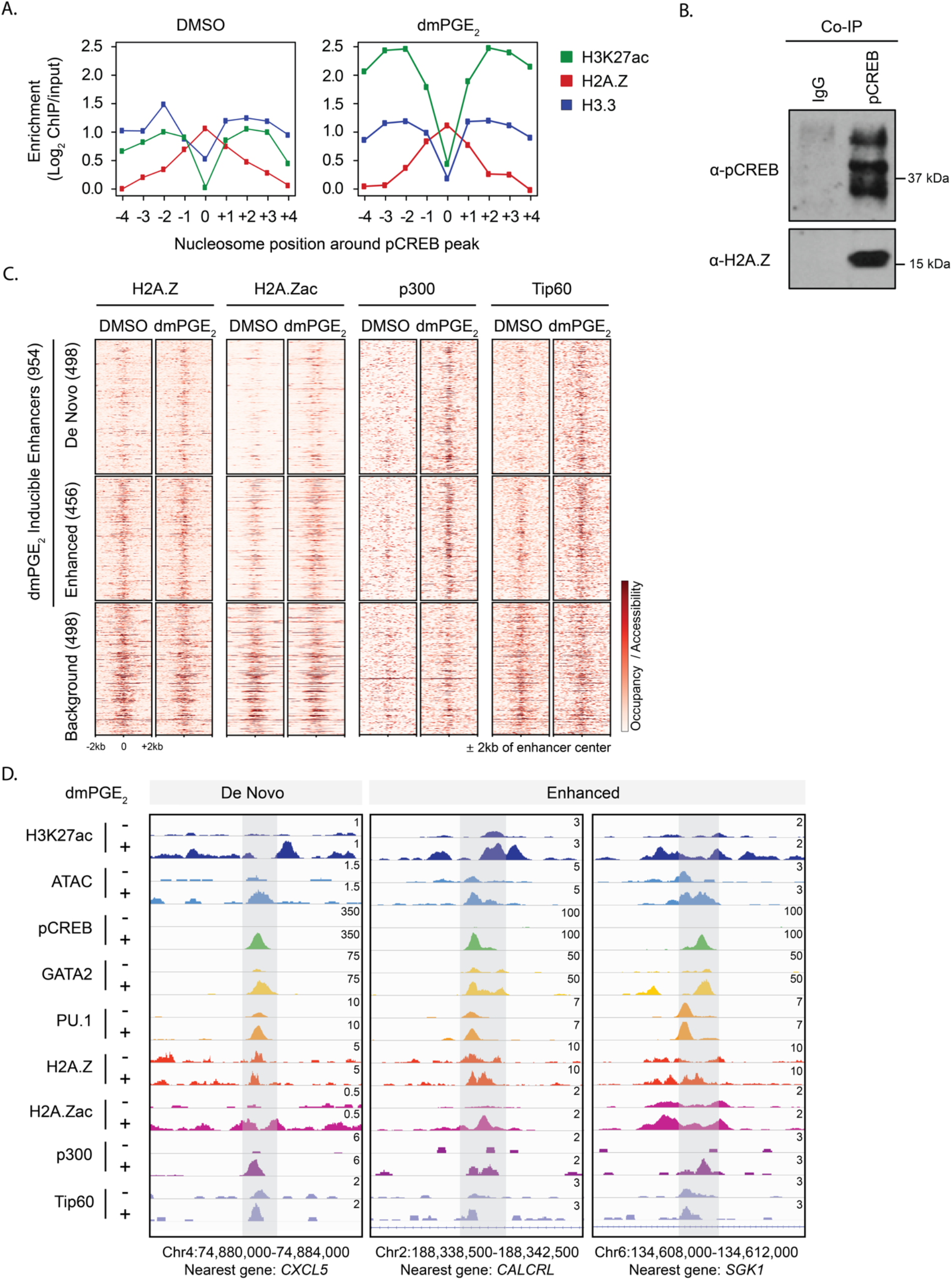
Modification of H2A.Z-variant accessible nucleosomes at stimuli-responsive enhancers by HATs p300 and Tip60. (A) Enrichment of histone variants at nucleosome positions surrounding pCREB peaks within enhancers before and after dmPGE_2_ stimulation. Position 0 indicates the nucleosome overlapping with pCREB peak centers. (B) Complex immunoprecipitation (Co-IP) showing that pCREB associated with H2A.Z in U937 myeloid leukemia cells (n = 3 biologically independent experiments). (C) Heat maps of H2A.Z, H2A.Zac, p300, and Tip60 binding at enhancers before and after dmPGE_2_ treatment. H3K27ac enriched regions identified by ChIP-Seq are classified as De Novo, Enhanced or Background enhancers according to the change in H3K27ac levels observed following dmPGE_2_ stimulation (n = 2 biologically independent experiments). (D) Enrichment of chromatin binding factors and transcription factors in response to dmPGE_2_ at 3 representative stimuli-response enhancers. Genomic location of presented window and enhancer nearest gene are indicated at the bottom of the panel. For all analysis presented here a randomly sampled set of background enhancers (486) was used.

H2A.Z is associated with both gene repression and activation^46^. This dual function is attributed to post-translational regulation of the histone variant. Different H2A.Z histone tail modifications recruit distinct interactors that mediate varying transcriptional outputs. H2A.Z acetylation is associated with active transcription and dynamically regulated in response to environmental signals^47,48^. Acetylation of H2A.Z occurs at active regulatory regions where it promotes nucleosomes destabilization and an open chromatin conformation. The histone acetyltransferases (HATs) p300 and Tip60 that acetylate H2A.Z are known interactors of pCREB^49,50^. Both chromatin remodelers hold important roles in hematopoietic stem cell fate^45,51–53^. Whereas Tip60 alone is not sufficient to acetylate H2A.Z, p300 can rapidly and effectively acetylate H2A.Z-containing nucleosomes on its own^54^. Hence, we assessed if acetylation of H2A.Z-variant nucleosomes at stimuli-responsive enhancers underlies increased MNase sensitivity and nucleosome accessibility after dmPGE_2_ treatment. We found specific H2A.Zac enrichment at stimuli-inducible enhancers whereas this was not observed in background enhancers (Figure 6C-D, Supplemental Figure 7D). We furthermore noted that dmPGE_2_ increased the abundance of both p300 and Tip60 at enhancers (Figure 6C-D, Supplemental Figure 7D). pCREB interacts with a variety of chromatin remodelers, including p300 and Tip60^49,54^. Binding patterns of two HATs at enhancers were highly similar to pCREB after dmPGE_2_ treatment (Figure 2B, Supplemental Figure 7D) and are suggestive of complex interaction at enhancers. Together, our data revealed that pCREB binds to H2A.Z-variant nucleosomes within enhancers following dmPGE_2_ stimulation. pCREB binding at stimuli-responsive enhancers is accompanied by nucleosome remodeling through H2A.Z acetylation, likely mediated through interaction of CREB with p300 and Tip60. H2A.Z acetylation at enhancers may underlie increased enhancer accessibility, allowing additional chromatin factors to engage and ultimately drive acute gene expression changes.

## Discussion

Prostaglandin E2 is an important regulator of HSPC homeostasis. The distinct molecular mechanisms through which PGE_2_ and its stable derivative dmPGE_2_ affect HSC function are critical to understand yet remain elusive. Here, we find that the TF CREB is a key player in the acute transcriptional response to dmPGE_2_ by binding to, and activating, distal regulatory elements. Specifically, we find that pCREB binds to H2A.Z-variant nucleosomes that are retained within stimuli-induced, active enhancers and is concomitant with acetylation of these histones. H2A.Z acetylation of enhancer nucleosomes increases local chromatin accessibility, which in turn may help to recruit and/or stabilize other HSPC specific TFs such as GATA2 to stimulate gene transcription.

CREB is a ubiquitously expressed nuclear, basic leucine zipper (bZIP) TF that regulates over 5,000 genes in the mammalian genome. This includes genes controlling proliferation, differentiation, and cell survival^55^. The TF is activated by a wide variety of environmental stimuli and is an important regulator of cellular responses to stress. Most studies focused on promoter-proximal effects of CREB binding. We showed that inducible gene expression is characterized by binding of pCREB at TSS-distal enhancers regions. Regulation of transcriptional responses through binding of CREB at enhancers has also been observed in pancreatic beta cells^56^. Our data suggests that CREB employs distinct mechanisms to regulate steady state versus inducible gene expression.

Genome wide assessment of the epigenetic landscape revealed that dmPGE_2_ works within the predetermined enhancer repertoire of HSPCs. dmPGE_2_ stimulation activates a set of pre-existing H3K4me1^+^ enhancers through chromatin reorganization. dmPGE_2_-responsive enhancers rapidly gain accessibility and TF binding. Our work complements the studies that showed that STFs localize to binding sites adjacent to master regulators^30,57^. Although we did not identify the surfacing of latent enhancers, *i.e*. genomic regulatory elements devoid of TFs and enhancer marks in unstimulated cells^27^, a 2-hour pulse of dmPGE_2_ may be too short to allow for partial reprogramming of the available cis-regulatory landscape. Latent enhancer activation may also be more associated with differentiated cells rather than stem and progenitor populations^27^.

We found stimuli-driven enrichment of the HSPC specific MTFs GATA2 and PU.1 at inducible enhancers, especially those that gained pCREB binding after 2 hours of treatment. Cofactor driven binding is common among non-pioneer TFs. It was only recently implied that interaction between TFs can enhance pioneer factor binding at previously sampled target sites^58–60^. Many studies have described the vital role of GATA2 in establishing the regulatory landscape for STFs in HSPCs^30,61^. GATA2 facilitates enhancer-promoter loop formation^62^, yet few have proposed signaling factors to play a role in GATA2 binding and recruitment to chromatin. We suggest that GATA2 occupancy may be directed and stabilized through cooperativity with STFs at stimuli-responsive enhancers. Our work supports a model where pioneer factor occupancy at specific subsets of enhancers is partially determined by engagement with signaling specific cofactors.

An open chromatin structure surrounding TF binding sites constitutes a prerequisite for transcriptional regulation. Our results show that inducible enhancers rapidly gain accessibility and affinity for STFs and MTFs. While open chromatin regions are presumed nucleosome depleted regions, we find that dmPGE_2_-responsive enhancers exhibit a high nucleosome occupancy both prior to and after stimulation of HSPCs. Moreover, retained enhancer nucleosomes are not prohibitive of STF binding. pCREB occupancy at enhancers overlaps with well-positioned nucleosomes, suggesting that pCREB can initiate chromatin engagement and access binding sites organized within a positioned nucleosome. Our work indicates that accessible nucleosomes at enhancers may facilitate cooperativity between STFs and MTFs to ensure rapid transcriptional induction. Retention of MNase accessible nucleosomes at regulatory elements was proposed to play a crucial role in hormone signaling and tissue specific gene activation^33,40^. Retained nucleosomes likely stabilize the interaction of TFs with DNA by facilitating interactions between TF-associated factors, such chromatin remodeling complexes and histone tails^40^.

Except for pioneer factors, most TFs are thought to be unable to bind nucleosomal DNA. Although CREB is not traditionally described as a pioneer factor, novel studies revealed the ability of CREB to open chromatin^63^. CREB was identified as a TF that displays an orientated, asymmetric, binding preference near the dyad axis of the nucleosome when engaging nucleosomal DNA^64,65^. We hypothesize that MNase accessible nucleosomes within stimuli-responsive enhancers enable cooperative TF binding for rapid gene activation.

Nucleosomes retained within stimuli-inducible enhancers were epigenetically pre-marked by the histone variant H2A.Z. Although the precise function of H2A.Z at enhancers at remains unclear, H2A.Z is an important regulator of enhancer activity in response to stimuli. H2A.Z-rich enhancers display higher chromatin accessibility and gene induction by promoting RNA polymerase II recruitment^42,46,66^. In contrast to hormone stimulation, which was found to increase H2A.Z incorporation at enhancer nucleosomes^67^, we found limited changes in H2A.Z distribution after dmPGE_2_ treatment. Instead, we noted that H2A.Z-variant nucleosomes undergo histone tail acetylation following dmPGE_2_ stimulation. Acetylated forms of H2A.Z are associated with an open chromatin conformation and directly regulate transcription of enhancer RNAs^68–70^. Our work implies that dmPGE_2_-inducible H2A.Z acetylation underlies increased low MNase sensitivity and enhanced nucleosomes accessibility at stimuli-responsive enhancers following dmPGE_2_ treatment. We found that changes in post-translational acetylation of H2A.Z at stimuli-responsive enhancers correlate with gene expression changes, indicating that H2A.Zac is a prerequisite for appropriate transcriptional induction following dmPGE_2_ stimulation.

Labile, H2A.Z-marked nucleosomes do not present an obstacle for pCREB binding, but may facilitate TF binding and enhancer activity. We suggest that a critical feature of pCREB is the recruitment of remodelers that opens the local nucleosome structure through H2A.Z acetylation in enhancers. pCREB interacts with a variety of chromatin remodelers. This includes the HATs p300 and Tip60, both known to interact with H2A.Z and catalyze its acetylation^49,54,71^. The recruitment of p300 to chromatin in a stimulus-dependent manner observed here is consistent with interactions between chromatin remodelers and other STFs^72–74^. Localization of p300 is concomitant to pCREB binding, suggesting pCREB-p300 complex formation at enhancers upon dmPGE_2_ stimulation. pCREB facilitated recruitment of Tip60 to enhancers complements previously described observations of Tip60 binding to a subset of enhancers and acetylating H2A.Z to promote expression of HSC genes^71–75^. The acetylation of H2A.Z-variant enhancer nucleosomes may create a chromatin environment permissive of enhancer activity and transcription.

In conclusion, this study reveals how specific genomic reorganization at a stimuli-responsive group of enhancers is directly translated into regulatory element activation and transcriptional induction. Our findings support a model where STFs and MTFs cooperate with nucleosomes to regulate activity of cis-regulatory elements that mediate adequate responses to environment signals. While the combination of cooperative lineage-specific MTF and inducible STF binding provides context and responsiveness to external stimuli, histone variant nucleosomes retained within inducible enhancers may facilitate TF binding by stabilizing chromatin complexes. Subsequent acetylation of histone variant nucleosomes by TF-associated nucleosome remodelers creates the accessible nucleosome landscape required at active transcriptional enhancers to ensure strong gene activation.

## Materials and methods

### Expansion of CD34^+^ cells

Human CD34^+^ hematopoietic stem and progenitor cells (HSPCs), isolated from peripheral blood of granulocyte colony-stimulating factor (G-CSF) mobilized healthy volunteers, were purchased from the Fred Hutchinson Cancer Research Center. The cells were maintained in suspension culture as previously described by Trompouki *et al*., 2011^30^. Briefly, the cells were expanded in StemSpan medium (Stem Cell Technologies Inc.) supplemented with StemSpan CC100 cytokine mix (Stem Cell Technologies Inc.) and 2% Penicillin-Streptomycin for a total of 6 days.

### Cell culture

U937 cells were maintained in suspension culture in RPMI-1640 supplemented with 10% (v/v) heat-inactivated fetal bovine serum, 1X Glutamax and 1% Penicillin-Streptomycin at 37° in a humidified atmosphere of 5% CO_2_.

### dmPGE_2_ treatment

16,16-dimethyl prostaglandin E2 was purchased reconstituted in DMSO from Cayman Chemicals (cat. #14750), aliquoted and stored in −80°C until use. Cells were counted, collected and resuspended in StemSpan medium with 2% PenStrep (CD34+ Cells) or RPMI with 1% Penicillin-Streptomycin, but in the absence of additional cytokines or growth factors. Cells were treated with dmPGE_2_ (Cayman chemicals) or DMSO (vehicle control) for 2 hours.

### qPCR analysis

RNA was extracted from using the RNeasy plus mini kit (Qiagen). cDNA synthesis was performed using the Superscript VILO (Invitrogen) and using equal amounts of starting RNA. The cDNA was analyzed with the Light Cycler 480 II SYBR green master mix (Applied Biosystems), and the QuantStudio 12K Flex (Applied Biosystems). All samples were prepared in triplicate. The PCR cycle conditions used are: (a) 95°C for 5 min, (b) [95°C for 10 sec, 54°C for 10 sec, 72°C for 15 sec] X 40 cycles. The analysis of Ct values were performed using 2^-ΔΔT method. The PCR primer-pairs used are:

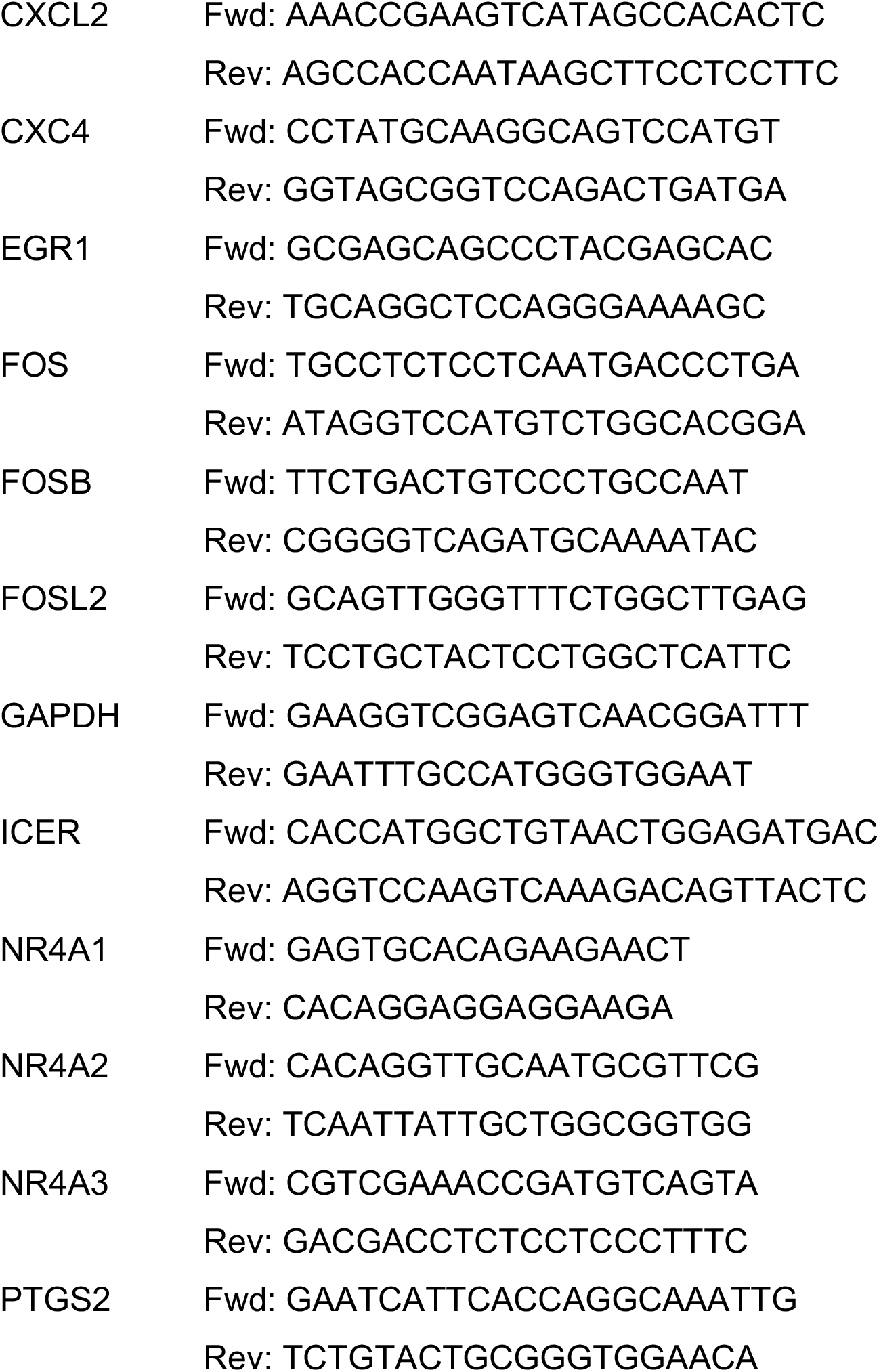

### Western blotting

Cells were treated for 2 hours, washed in 1X PBS, and collected in RIPA buffer with protease and phosphatase inhibitors. Samples were run on acrylamide gel and transferred onto a nitrocellulose membrane. Membrane was blocked for one hour in 5% of milk or TBS-T and incubated overnight at 4°C with anti-vinculin (Abcam 73412), anti-CREB (Santa Cruz SC-186), anti-pCREB (Ser133, Cell Signalling #9198), anti-H2A.Z (Abcam 4174), or anti-IgG (Cell Signalling #2729). The next day, membranes were washed, incubated with HRP-conjugated secondary antibodies for 1 hour at room temperature, and developed with SuperSignal West Pico Plus Chemiluminescent substrate.

### Co-immunoprecipitation (Co-IP)

Co-IP was performed as previously described by Santoriello *et al*., 2020^76^. Cells were washed twice with 1X PBS. Nuclei were isolated with 0.05% Triton in PBS and lysed in nuclei lysis buffer (20mM Hepes-KOH pH7.9, 25% glycerol, 420mM NaCl, 1.5mM MgCl2, 0.2mM EDTA, 0.5mM DTT). 500ug of protein extracts was used and the salt concentration was diluted from 420mM to 150mM NaCl using 20mM Hepes-KOH pH7.9, 20% glycerol, 0.25mM EDTA, 0.05% NP-40. Cell lysates were pre-cleared with non-antibody bounds beads for 1 hour at 4°C. Antibody bounds protein G Dynabeads were added to pre-cleared lysate and samples incubated overnight at 4°C. Antibodies used: IgG (Cell Signalling #2729), phospho-CREB (Ser133, Cell Signalling #9198). Protein-bead complexes were then washed 5 times with wash buffer (20mM Hepes-KOH pH7.9, 10% Glycerol, 150mM NaCl, 1.5 MgCl2, 0.2mM EDTA, 0.5mM DTT) and beads were boiled in 50μL Laemmli Buffer for 15min at 95°C to elute proteins. Subsequently, samples were subjected to western blot.

### RNA-Seq

RNA from one million cells was isolated using the RNeasy plus mini kit (Qiagen #74134). 5μg of RNA was subjected to ribosomal and mitochondrial RNA depletion using the RiboZero Gold kit (Human/Mouse/Rat, Epicentre #MRZG12324) according to manufacturer’s instructions. The ribo-zero treated RNA wa*s* used to create multiplexed RNA-Seq libraries using the NEBNext Ultra RNA Library Prep Kit (Illumina E7530) according to the manufacturer’s instructions. Briefly 500pg of ribozero treated RNA was fragmented and used to produce cDNA libraries using the NEBnext Ultra RNA library prep kit (NEB, E7530S) according to the manufacturer’s protocol. Purified double-stranded cDNA underwent end-repair and dA-tailing reactions following manufacturer’s reagents and reaction conditions. The obtained DNAs were used for Adaptor Ligation using adaptors and enzymes provided in NEBNext Multiplex Oligos for Illumina (NEB #E7335) and following recommended reaction conditions. Eluted DNA was enriched with PCR reaction using Fusion High-Fidelity PCR Master Mix kit (NEB, M0531S) and specific index primers supplied in NEBNext Multiplex Oligo Kit for Illumina (Index Primer Set 1, NEB, E7335L). Conditions for PCR used are as follows: 98°C, 30 sec; [98°C, 10 sec; 65°C, 30 sec; 72°C, 30 sec] X 15 cycles; 72°C, 5 min; hold at 4°C. PCR reaction mix was purified using Agencourt AMPure XP beads (1X of reaction volume). Libraries were eluted in 20μl elution buffer. All the libraries went through quality control analysis using an Agilent Bioanalyzer and subjected to next-generation sequencing using Illumina Hiseq 2500 platform. Quality control of RNA-Seq datasets was performed by FastQC and Cutadapt to remove adaptor sequences and low-quality regions. The high-quality reads were aligned to UCSC hg19 for human using Tophat 2.0.11 without novel splicing form calls. Transcript abundance and differential expression were calculated with Cufflinks 2.2.1. FPKM values were used to normalize and quantify each transcript.

### ChIP-Seq

For ChIP-Seq experiments the following antibodies were used: H3K27ac (Abcam ab4729), H3K4me1 (Abcam ab8895), H3K27me3, (Abcam ab195477), pCREB (Ser133, Cell Signalling #9198), GATA2 (Santa Cruz sc9008X), H2A.Z (Abcam ab4174), H2A.Zac (Abcam ab18262), H2B (Abcam ab1790), H3.3 (Milipore #09-838), H4 (Abcam ab7311), p300 (Millipore #05-257), and Tip60 (generous gift from Bruno Amati). ChIP experiments were performed as previously described by Trompouki *et al*., 2011^30^. Briefly, 20 million cells were crosslinked by the addition of 1/10 volume 11% fresh formaldehyde for 10 min at room temperature. The crosslinking was quenched by the addition of 1/20 volume 2.5M glycine for 5 minutes. Cells were washed twice with ice-cold PBS. Cells were lysed in 10mL of Lysis buffer 1 (50mM HEPES-KOH, pH 7.5, 140mM NaCl, 1mM EDTA, 10% glycerol, 0.5% NP-40, 0.25% Triton X-100, plus protease and phosphatase inhibitors) for 10 min at 4°C. After centrifugation, cells were resuspended in 10 mL of Lysis buffer 2 (10mM Tris-HCl, pH 8.0, 200mM NaCl, 1mM EDTA, 0.5mM EGTA, plus protease and phosphatase inhibitors) for 10 min at room temperature. Cells were pelleted and resuspended in 3mL of sonication buffer (10mM Tris-HCl, pH 8.0, 100mM NaCl, 1mM EDTA, 0.5mM EGTA, 0.1% Na-Deoxycholate, 0.05% N-lauroylsarcosine, plus protease and phosphatase Inhibitors) and sonicated in a Bioruptor sonicator for 36 cycles of 30 sec each followed by a 1min resting interval. Samples were centrifuged for 10min at 18,000g and 1% Triton-X was added to the supernatant. Prior to the immunoprecipitation, 50mL of protein G beads (Invitrogen 100-04D) for each reaction were washed twice with PBS, 0.5% BSA twice. Finally, the beads were resuspended in 250μL of PBS, 0.5%BSA and 5μg of each antibody. Beads were rotated for at least 6 hours at 4°C and then washed twice with PBS with 0.5% BSA. Cell lysates were added to the beads and incubated at 4°C overnight. Beads were washed 1x with (20mM Tris-HCl. pH 8, 150mM NaCl, 2mM EDTA, 0.1% SDS, 1% Triton X-100), 1x with (20mM Tris-HCl, pH 8, 500mM NaCl, 2mM EDTA, 0.1% SDS, 1% Triton X-100), 1x with (10mM Tris-HCl, pH 8, 250nM LiCl, 2mM EDTA, 1% NP4-0) and 1x with TE and finally resuspended in 200μL elution buffer (50mM Tris-Hcl, pH 8.0, 10mM EDTA, 0.5%–1% SDS). 50μL of cell lysates prior to addition to the beads was kept as input. Crosslinking was reversed by incubating samples at 65°C for at least 6 hours. Afterwards the cells were treated with RNase and proteinase K and the DNA was extracted by Phenol/Chloroform extraction.

ChIP-Seq libraries were prepared using the following protocol. End repair of immunoprecipitated DNA was performed using the End-It End-Repair kit (Epicentre, ER81050) and incubating the samples at 25°C for 45 min. End-repaired DNA was purified using AMPure XP Beads (1.8X of the reaction volume) (Agencourt AMPure XP – PCR purification Beads, BeckmanCoulter, A63881) and separating beads using DynaMag-96 Side Skirted Magnet (Life Technologies, 12027). A-tails were added to the end-repaired DNA using NEB Klenow Fragment Enzyme (3’-5’ exo, M0212L), 1X NEB buffer 2 and 0.2mM dATP (Invitrogen, 18252-015) and by incubating the reaction mix at 37μC for 30 min. A-tailed DNA was cleaned up using AMPure beads (1.8X of reaction volume). Subsequently, cleaned up A-tailed DNA went through Adaptor ligation reaction using Quick Ligation Kit (NEB, M2200L) following manufacturer’s protocol. Adaptor-ligated DNA was first cleaned up using AMPure beads (1.8X of reaction volume), eluted in 100μl and then size-selected using AMPure beads (0.9X of the final supernatant volume, 90μl). Adaptor-ligated DNA fragments of proper size were enriched with PCR reaction using Fusion High-Fidelity PCR Master Mix kit (NEB, M0531S) and specific index primers supplied in NEBNext Multiplex Oligo Kit for Illumina (Index Primer Set 1, NEB, E7335L). Conditions for PCR used are as follows: 98°C, 30 sec; [98°C, 10 sec; 65°C, 30 sec; 72°C, 30 sec] X 15 to 18 cycles; 72°C, 5 min; hold at 4°C. PCR enriched fragments were cleaned up using AMPure beads (1X of reaction volume). Libraries were eluted in 20μl elution buffer. All the libraries went through quality control analysis using an Agilent Bioanalyzer and subjected to next-generation sequencing using Illumina Hiseq 2500 platform. All the libraries went through quality control analysis using an Agilent Bioanalyzer and subjected to next-generation sequencing using Illumina Hiseq 2500 platform. All ChIP-Seq datasets were aligned to UCSC build version hg19 of the human genome using Bowtie2 (version 2.2.1; Langmead *et al*., 2012^77^) with the following parameters: -end-to-end, -N0, -L20. We used the MACS2 version 2.1.0 (Zhang *et al*., 2008^78^) peak-finding algorithm to identify regions of ChIP-Seq peaks, with a q-value threshold of enrichment of 0.05 and false discovery rate of < 0.01 for all datasets. The genome-wide occupancy profile figures were generated by deeptools (Ramirez *et al*., 2016^79^) using the reference-point mode and the scale-regions mode. The genomic distribution of peaks was plotted using the ChIPSeeker R package, annotatePeak to assign peaks to a genomic annotation, which includes whether a peak is in the TSS, Exon, 5’ UTR, 3’ UTR, Intronic or Intergenic. The genome annotation is from the R-bioconductor annotation packages. Heat maps of the ChIP-Seq binding were generated using the input normalized results of the MACS peak calling output. The outputted bedGraph files were converted to BigWig files. Those files were then processed using computeMatrix and plotHeatmap tools within the deeptools 3.0 package. Enhancers were assigned to genes using GREAT to the nearest genes within 15kb of peaks.

### Defining Enhancer Categories

We used the SPP package to call clusters of H3K27ac enrichment, normalized to input, from ChIP-Seq data (Kharchenko *et al*., 2008^80^). Regions within 500bp of each other were merged and only regions reproduced between two independent biological H3K27ac ChIP-Seq replicates (in either sample dmPGE_2_ or DMSO) were included for further analysis. Enhancers were defined as TSS distal H3K27ac regions that are ≥ 1kb in length and located ≥ 2kb away from TSS. This yielded a total of 25,998 H3K27ac enriched ChIP-Seq regions, here named enhancers. P-value was computed using paired t-test between dmPGE_2_ and DMSO on enrichment values for every region.

Called enhancers were classified based on the three following criteria and using cut-off described in the table below: (1) H3K27ac enrichment in each replicate in each condition, (2) delta H3K27ac enrichment upon stimulation, (3) p-value of Δ H3K27ac enrichment. All regions not classified as not meeting above mentioned criteria were classified as ‘Background’ enhancers.

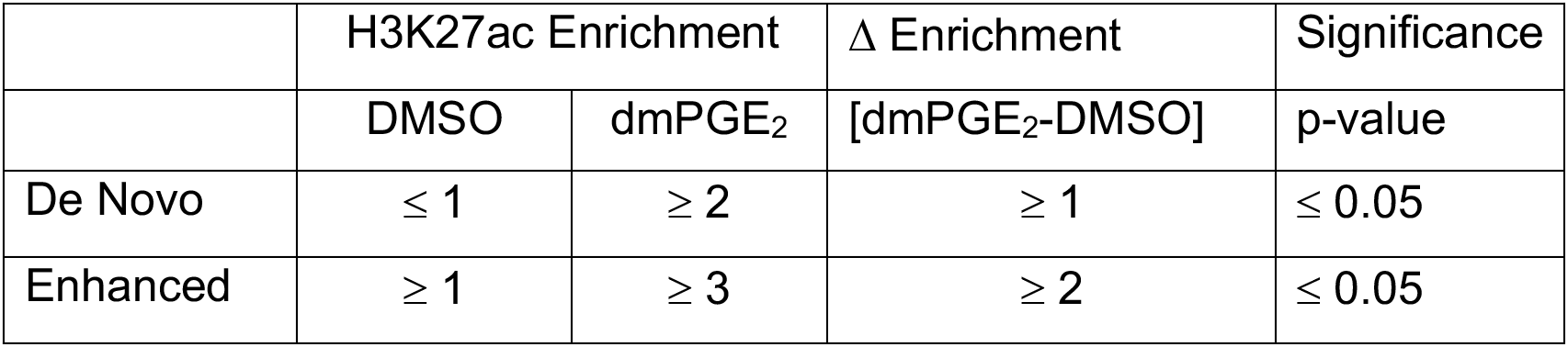

| | H3K27ac Enrichment | | $\Delta$ Enrichment | Significance |
| --- | --- | --- | --- | --- |
|  | DMSO | dmPGE<sub>2</sub> | [dmPGE<sub>2</sub>-DMSO] | p-value |
| De Novo | $\leq 1$ | $\geq 2$ | $\geq 1$ | $\leq 0.05$ |
| Enhanced | $\geq 1$ | $\geq 3$ | $\geq 2$ | $\leq 0.05$ |

### ATAC-Seq

50.000 cells per condition were harvested by spinning at 500g for 5 min, 4°C. Cells were washed once with 50μl of cold 1X PBS and spun down at 500g for 5 min, 4°C. After discarding supernatant, cells were lysed using 50μl cold lysis buffer (10mM Tris-HCl pH 7.4, 10mM NaCl, 3 mM MgCl2, 0.1% IGEPAL) and spun down immediately at 500g for 10 min at 4°C. Then the cells were precipitated and kept on ice and subsequently resuspended in 25μl 2X Tagment DNA Buffer (Illumina Nextera kit), 2.5μl Transposase enzyme (Illumina Nextera kit, 15028252) and 22.5μl Nuclease-free water in a total of 50uL reaction for 1 hour at 37°C. DNA was then purified using Qiagen MinElute PCR purification kit (28004) in a final volume of 10μl. Libraries were constructed according to Illumina protocol using the DNA treated with transposase, NEB PCR master mix, Sybr green, universal and library-specific Nextera index primers. The first round of PCR was performed under the following conditions: 72°C, 5 min; 98°C, 30 sec; [98°C, 10 sec; 63°C, 30 sec; 72°C, 1 min] X 5 cycles; hold at 4°C. Reactions were kept on ice and using a 5μL reaction aliquot, the appropriate number of additional cycles required for further amplification was determined in a side qPCR reaction: 98°C, 30 sec; [98°C, 10 sec; 63°C, 30 sec; 72°C, 1 min] X 20 cycles; hold at 4°C. Upon determining the additional number of PCR cycles required further for each sample, library amplification was conducted using the following conditions: 98°C, 30 sec; [98°C, 10 sec; 63°C, 30 sec; 72°C, 1 min] X appropriate number of cycles; hold at 4°C. Libraries prepared went through quality control analysis using an Agilent Bioanalyzer and then subjected to next generation sequencing using Illumina Hiseq 2500 platform. We used the MACS2 version 2.1.0 (Zhang *et al*., 2008^78^) peak-finding algorithm to identify regions of ATAC-Seq peaks, with the following parameter -- nomodel --shift −100 --extsize 200. A q-value threshold of enrichment of 0.05 was used for all datasets.

### MNase-Seq

CD34^+^ HSPC cells were crosslinked by the addition of 1/10 volume 11% fresh formaldehyde for 10 min at room temperature. The crosslinking was quenched by the addition of 1/20 volume 2.5M glycine for 5 minutes. Cells were washed twice with ice-cold PBS. For MNase digestion, the nuclei pellet was resuspended in MNase digestion buffer (50mM Tris, pH 7.4, 25mM KCl, 4mM MgCl2, 1mM CaCl2, 12.5% Glycerol and COMPLETE protease inhibitors (Roche)). Digestion took place with 10^6^ cells per titration point in a volume of 500μl MNase digestion buffer. Either 1, 2, 4, 8, 16, 32, 64, 128 or 256 units of MNase (Worthington Biochemical) were added to pre-warmed nuclei and incubated at 25 °C for 15 min. Digestion was halted with 25mM EDTA/EGTA and 0.5% SDS and 125mM NaCl was added to the samples. Digestions were incubated with RNase (Roche) for 1 hour at 37 °C, with proteinase K (Roche) for 1 hour at 55 °C, and cross-link reversal was performed at 65 °C for 16 hours. DNA was purified by Phenol/Chloroform extraction and ethanol precipitation. MNase digestion was evaluated on a 2% agarose gel and fragments from four MNase concentrations representing 10%, 25%, 50% and 75% mono-nucleosomal fragments were individually prepared for next generation sequencing. Ampure SPRI beads (Beckman Coulter) were used in a double size selection with ratios of 0.7X and 1.7X to obtain a range of fragment sizes from ~100 to 1,000bp. DNA was eluted from the beads and used as input into the library preparation protocol. DNA libraries were prepared for each individual titration point using the NEBNext Ultra II DNA Library Prep Kit for Illumina (E7370, New England Biolabs) and barcoded using NEBNext Multiplex Oligos for Illumina (Index Primers Set 1 & 2; New England Biolabs). Number of PCR cycles was calculated using a real-time qPCR-based approach (Lion *et al*., 2020^81^). Libraries prepared went through quality control analysis using an Agilent Bioanalyzer. Four barcoded titration libraries were pooled in one sample, and paired-end sequencing on an Illumina Hiseq 2500 instrument was performed. Three biological replicated were sequenced. The sequenced paired-end reads were mapped to hg19 using Bowtie aligner v. 0.12.9. Only uniquely mapped reads with no more than two mismatches were retained. The reads with the insert sizes <50 bp or >500 bp were filtered out. Genomic positions with the numbers of mapped tags above the significance threshold of *Z*-score=7 were identified as anomalous, and the tags mapped to such positions were discarded. Read frequencies were computed in 300bp non-overlapping bins for each titration point independently. The read frequencies were normalized by the corresponding library sizes to represent values per one million of mapped reads. Nucleosome occupancy analysis was carried out as previously described by Mieczkowski *et al*., 2016^32^.

## Acknowledgments

We thank J.M. Ordovas-Montanes for critical reading of our manuscript; A. Choudhuri, R. E. Kingston and the members of the Zon laboratory for discussions on the project. A.S. was supported by a Boehringer Ingelheim PhD Fellowship. A.S. was supported by a Boehringer Ingelheim PhD Fellowship. This work was supported by the following grants from L.I.Z.: R01 HL04880, P015PO1HL32262-32, 5P30 DK49216, 5R01 DK53298, 5U01 HL10001-05, R24 DK092760, 1R24OD017870-01.

## Author Contributions

A.S., E.M.F, and M.E.M., performed experiments. M.P., B.M. and S.Y. performed bioinformatics analysis of next generation sequencing data. A.C., R.E.K., M.Y.T, and H. C. provided insights on data analysis and interpretation. A.S. and L.I.Z. managed the study and wrote the manuscript.

