## Supplemental Figures for "Stimulation-responsive enhancers regulate inflammatory gene activation through retention and modification of H2A.Z-variant accessible nucleosomes"

### Supplemental Figure 1

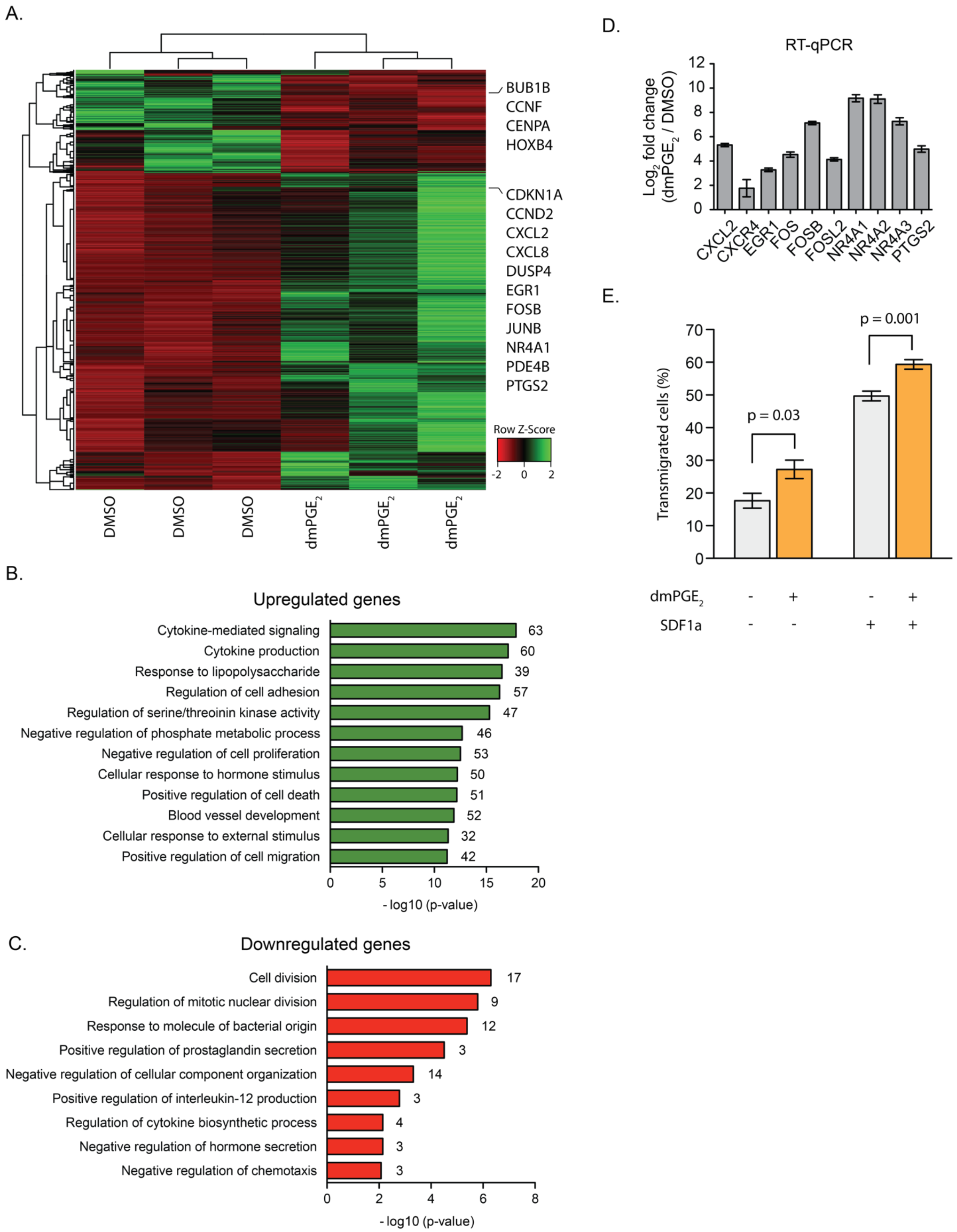

**Supplemental Figure 1. dmPGE$_2$ induces acute transcriptional responses in HSPCs.** (A) Hierarchical clustering heatmap of FPKM values from differentially expressed genes (687) 2 hours post-treatment. DEG criteria: FPKM ≥ 1 after treatment; fold change ≥ 1.5 or ≤ 0.67 (n = 3 biologically independent experiments). (B, C) Gene Ontology (GO) term enrichment analysis of genes upregulated (C, in green) and down-regulated (D, in red) in CD34$^+$ HSPCs 2 hours post dmPGE$_2$ treatment. The number of genes associated with each GO term are shown at the end of the bar within the graph. P-values were calculates using hypergeometric test and Benjamini-Hochberg correction. (D) RT-qPCR in CD34$^+$ HSPCs of genes identified as differentially expressed by RNA-Seq (n = 3 biologically independent experiments; mean values ± SEM). (E) CD34$^+$ HSPCs were exposed to dmPGE$_2$ or DMSO for 2h cells after which the stimuli were washed out. Cells were then placed in the top chamber of the transwell system, with or without recombinant human SDF-1α in the bottom chamber. After 24 hours, cells migration to the bottom chamber was quantified as percentage of total cells seeded. (n = 3 biologically independent experiments; mean values ± SEM).

### Supplemental Figure 2

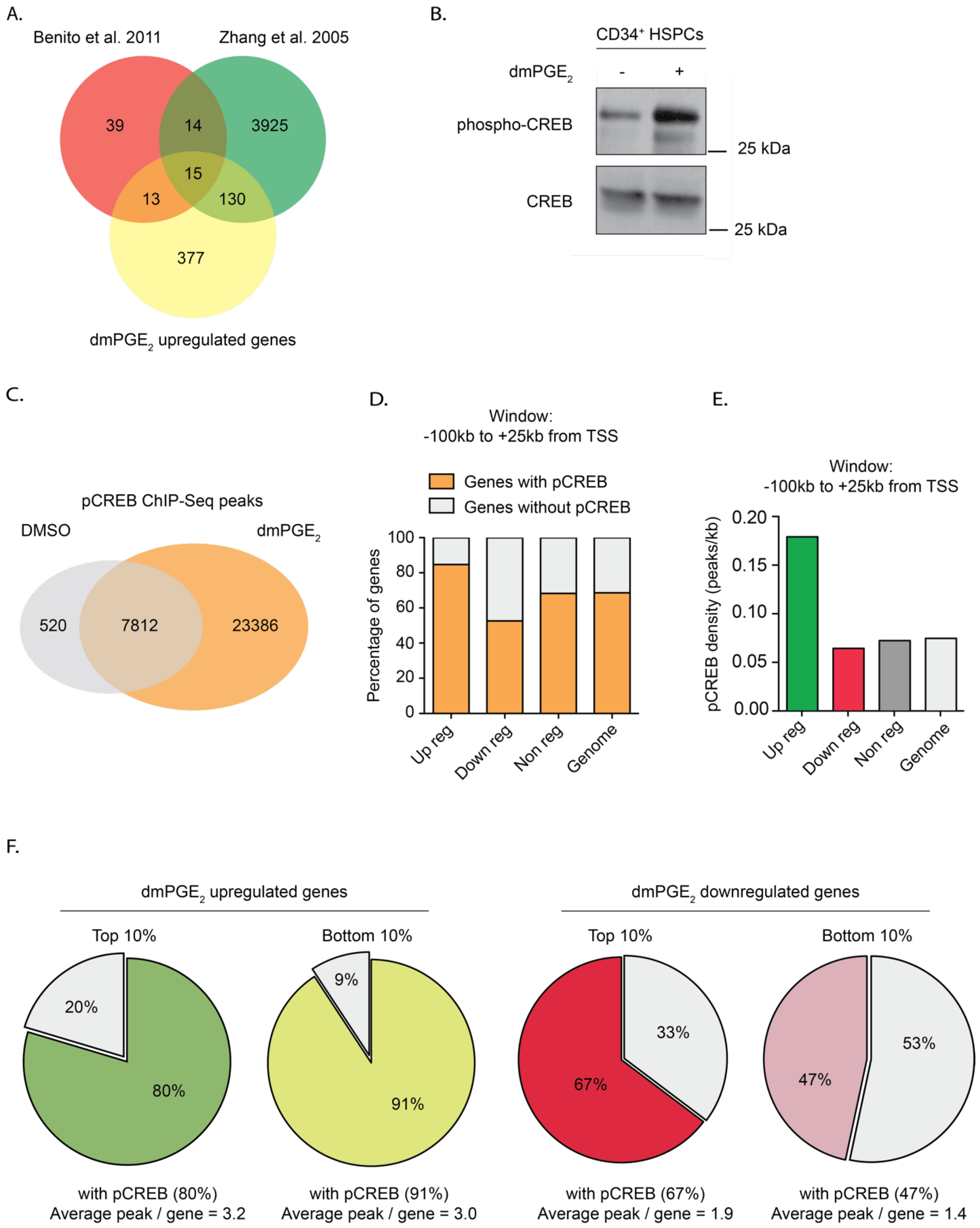

**Supplemental Figure 2. pCREB binds near differentially expressed genes.** (A) Venn diagram showing overlap between upregulated genes (535) and previously identified CREB target genes. (B) Western blot analysis for phospho-CREB in $CD34^+$ HSPCs stimulated with vehicle control (DMSO) or $dmPGE_2$ for 2 hours. Total CREB protein was used as loading control. (C) Venn diagram showing overlap between pCREB peaks present in DMSO after $dmPGE_2$ stimulation, as identified by ChIP-Seq. (D) Number of genes containing at least one pCREB peak in the proximity after $dmPGE_2$ stimulation. pCREB peaks were assigned to a gene when located within a window from -100kb upstream of the transcription start site (TSS) to +25kb downstream of the TTS was considered (n = 2 biologically independent ChIP-Seq experiments). (E) Correlation between pCREB binding and gene expression in response to $dmPGE_2$. pCREB density was calculated by dividing the total number of pCREB peaks associated to each gene category (up-, down-, and nonregulated genes) by the total amount of base pairs that this category occupies in the genome. pCREB peaks were assigned to a gene when located from +100kb upstream of the TSS to +25kb downstream of the TTS. Peak density in the genome was calculated by considering random distribution of pCREB sites in the whole genome. (F) pCREB in $dmPGE_2$-response genes. Top and bottom 10% correspond to the 10% most upregulated and downregulated genes, respectively. pCREB peaks were assigned to a gene when located within a window from -5kb upstream of the transcription start site (TSS) to +5kb downstream of the TTS was considered (n = 2 biologically independent ChIP-Seq experiments).

### Supplemental Figure 3

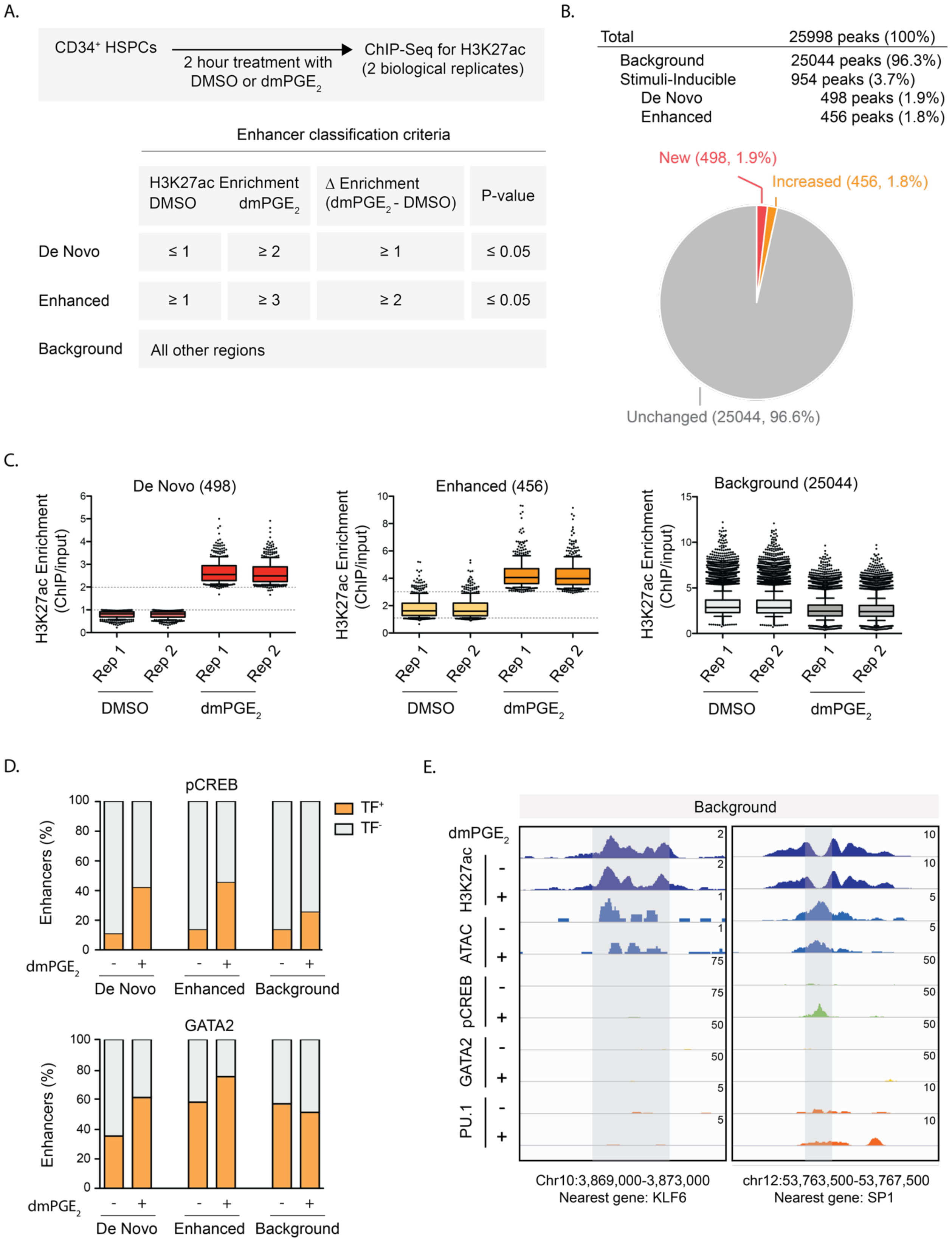

**Supplemental Figure 3. Identification of stimuli-inducible enhancers in HSPCs.** (A) Experimental set up and enhancer classification criteria. (B) Number of identified enhancer regions and their distribution into the different categories based on two independent replicate experiments. (C) H3K27ac enrichment levels at enhancers within each indicated category for 2 biologically independent replicate (rep) ChIP-Seq experiments. Dotted lines indicate cutoff values used for enhancer classifications. (D) Number of enhancers as percentage of total within each category containing enrichment for pCREB (upper panel) and GATA2 (lower panel). (E) Enrichment of histone mark, ATAC accessibility and transcription factor binding in response to dm$PGE_2$ at representative 2 background enhancers. Genomic location of presented window and nearest gene are indicated at the bottom of the panel.

### Supplemental Figure 4

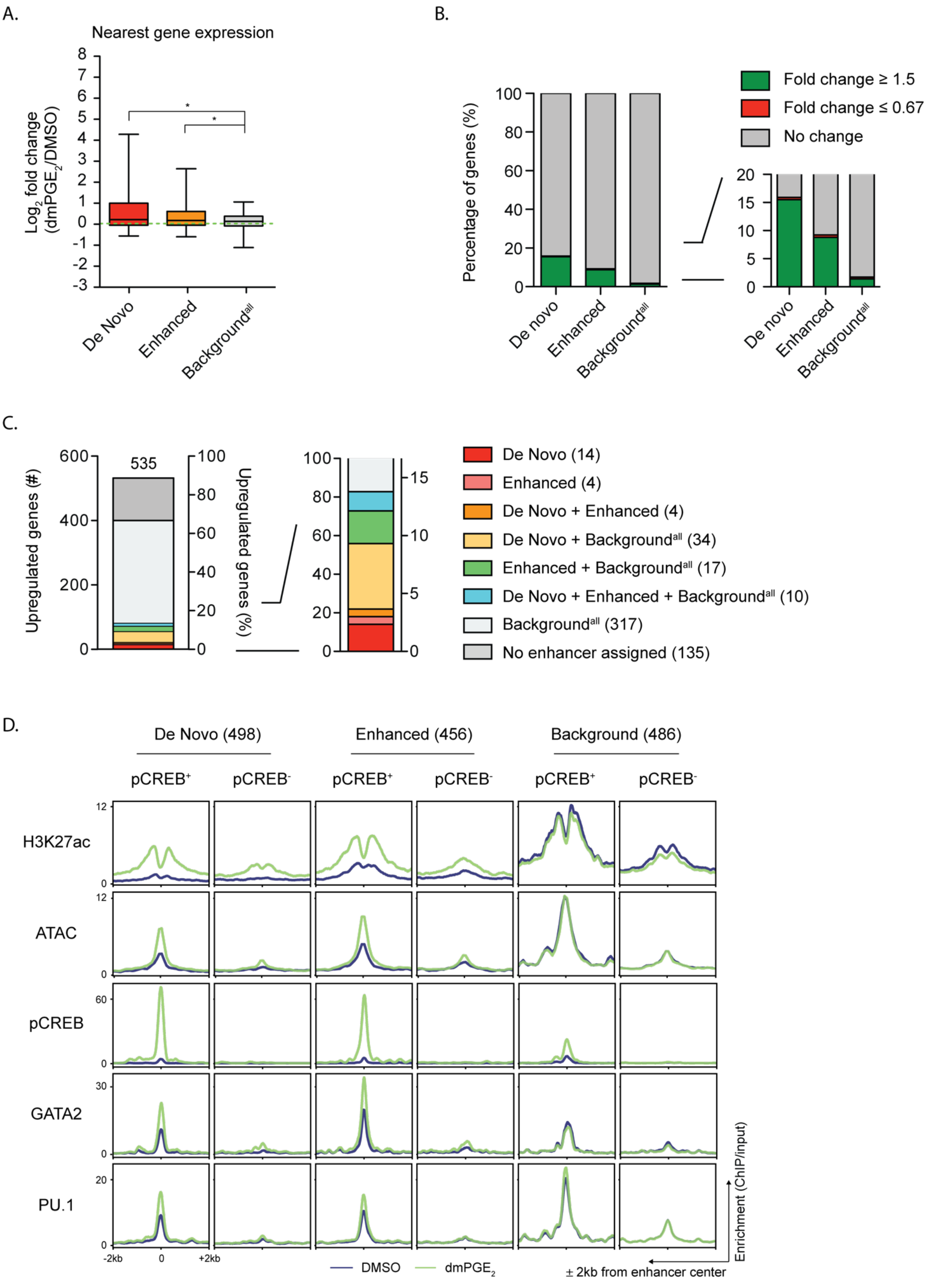

**Supplemental Figure 4. Stimuli-responsive enhancers mediate gene expression changes.** (A) Gene expression changes of genes associated with stimuli-responsive and background enhancers. Enhancers were assigned to an individual nearest gene. Only genes with a mapped TSS within 15kb of an enhancer were considered. Box plots shows median, 25th and 75th percentiles, whiskers are from 5th and 95th percentiles. (B) Percentages of enhancer nearest genes with fold changes in expression ≥ 1.5-fold or ≤ 0.67-fold for each enhancer category. (C) Upregulated genes with a fold change in expression ≥ 1.5 (535) and their associated enhancers. For all analysis presented in A, B, and C the entire set of background enhancers (25,044) was used. (D) Average enrichment profile of H3K27ac, ATAC accessibility and transcription factors around enhancers before and after $dmPGE_2$ treatment. De Novo, Enhanced or Background enhancers were subset based on the presence or absence of pCREB after $dmPGE_2$. A randomly sampled, comparable number of background enhancers (486) is shown. * = $p < 0.0001$

### Supplemental Figure 5

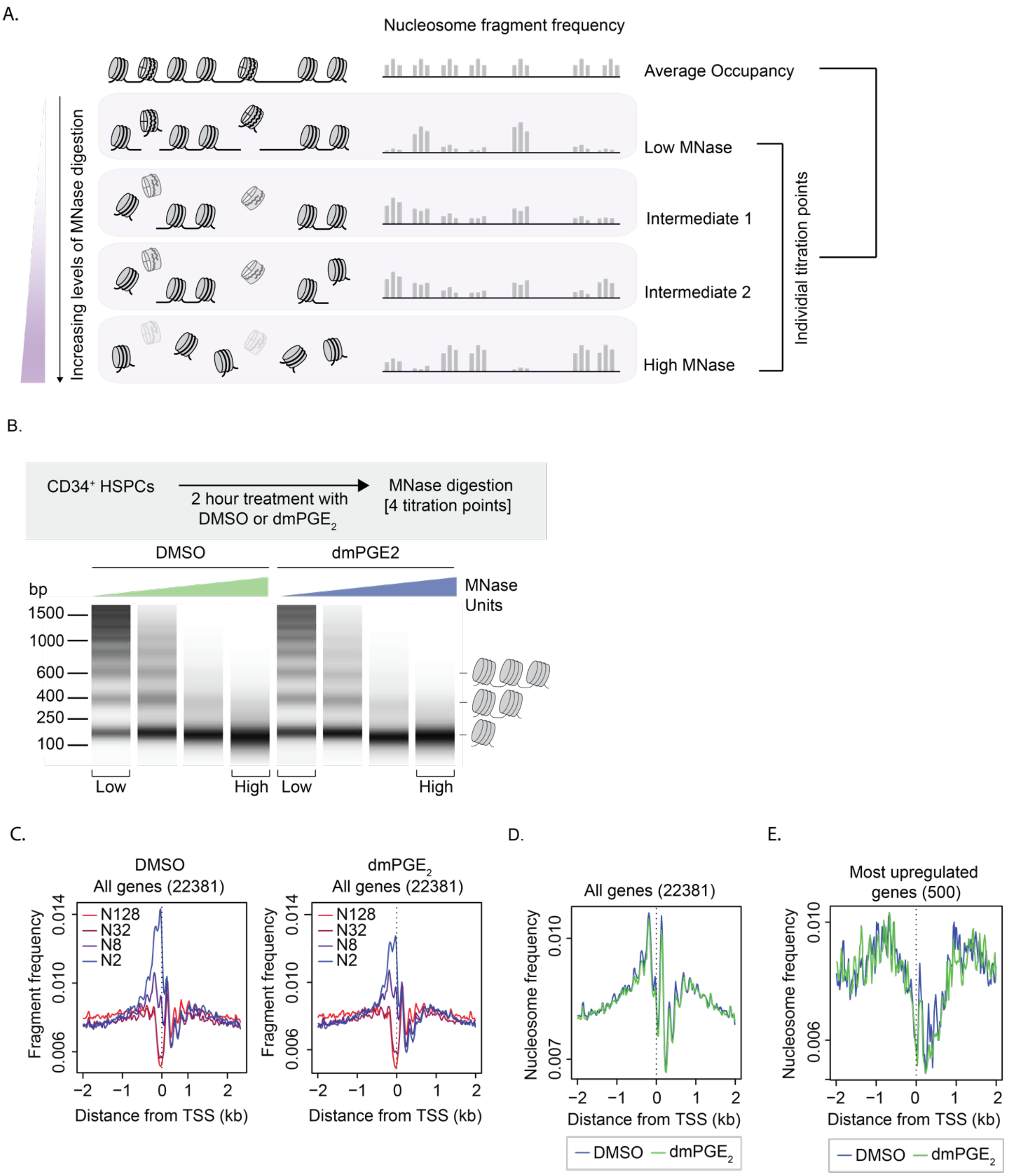

**Supplemental Figure 5. MNase-sequencing in DMSO and $dmPGE_2$ treated HSPCs.** (A) Schematic representation of MNase-Seq using 4 titration point. Average nucleosome occupancy is determined through pooled analysis of 4 individual MNase digestion levels, as indicated (n = 3 independent biological experiments). (B) Capillary electrophoresis of digestion products from a typical MNase titration experiment. Cells are stimulated for 2 hours with $dmPGE_2$ or vehicle control (DMSO) after which MNase digestion was performed. (C) MNase-Seq profiles around TSS (transcription start sites) of all genes. Colors indicates MNase concentration levels (2, 8, 32 and 128 Units of MNase), with blue corresponding to the lowest concentration and red corresponding to the highest. (D) The average nucleosome profile at the TSS of all genes in DMSO treated (blue) and $dmPGE_2$ (green) treated HSPCs, as determined from 4 individual MNase titration point per experimental condition. (E) The average nucleosome profile at the TSS of the 500 most upregulated in DMSO treated (blue) and $dmPGE_2$ (green) treated HSPCs, as determined from 4 individual MNase titration point per experimental condition.

**Supplemental Figure 6**

A.

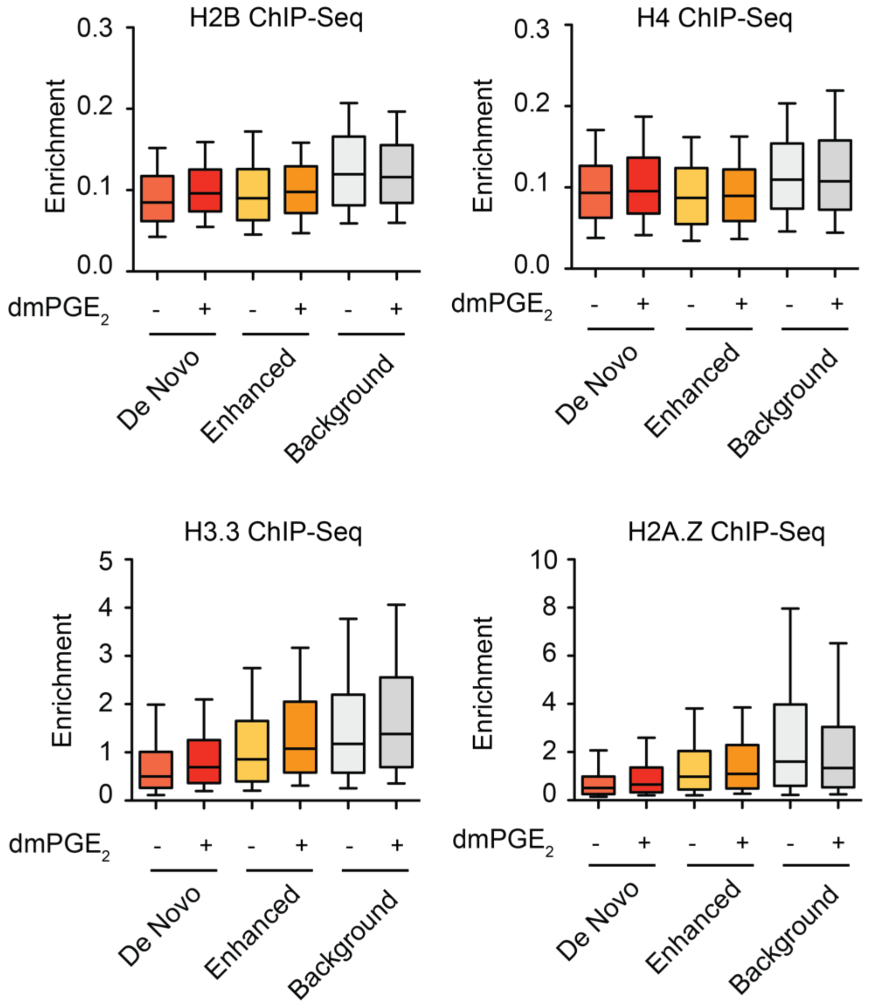


B.

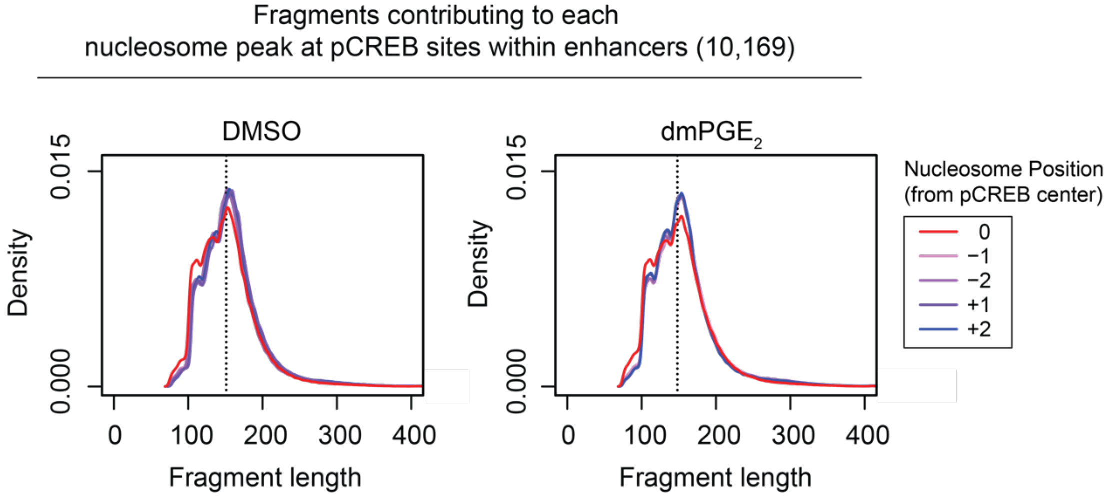

**Supplemental Figure 6. Histone enrichment and MNase fragment size at enhancers.** (A) Box and whisker plots show H2B, H3, H2A.Z and H3.3 ChIP-seq signal enrichment at the central 500bp of stimuli-responsive and background enhancers, in DMSO and $dmPGE_2$ treated HSPCs. Box plots shows median, $25^{th}$ and $75^{th}$ percentiles, whiskers are from $10^{th}$ and $90^{th}$ percentiles. For analysis presented here, a randomly sampled comparable number of background enhancers (486) is used. (B) Size distribution of DNA fragment reads mapped to the corresponding nucleosome position displayed. Sequencing libraries prepared from MNase-generated fragments were subjected to paired-end sequencing, and the sizes of the fragments were inferred from the positions of the mapped ends. Nucleosome position 0 indicates the nucleosome overlapping with pCREB peak centers.

### Supplemental Figure 7

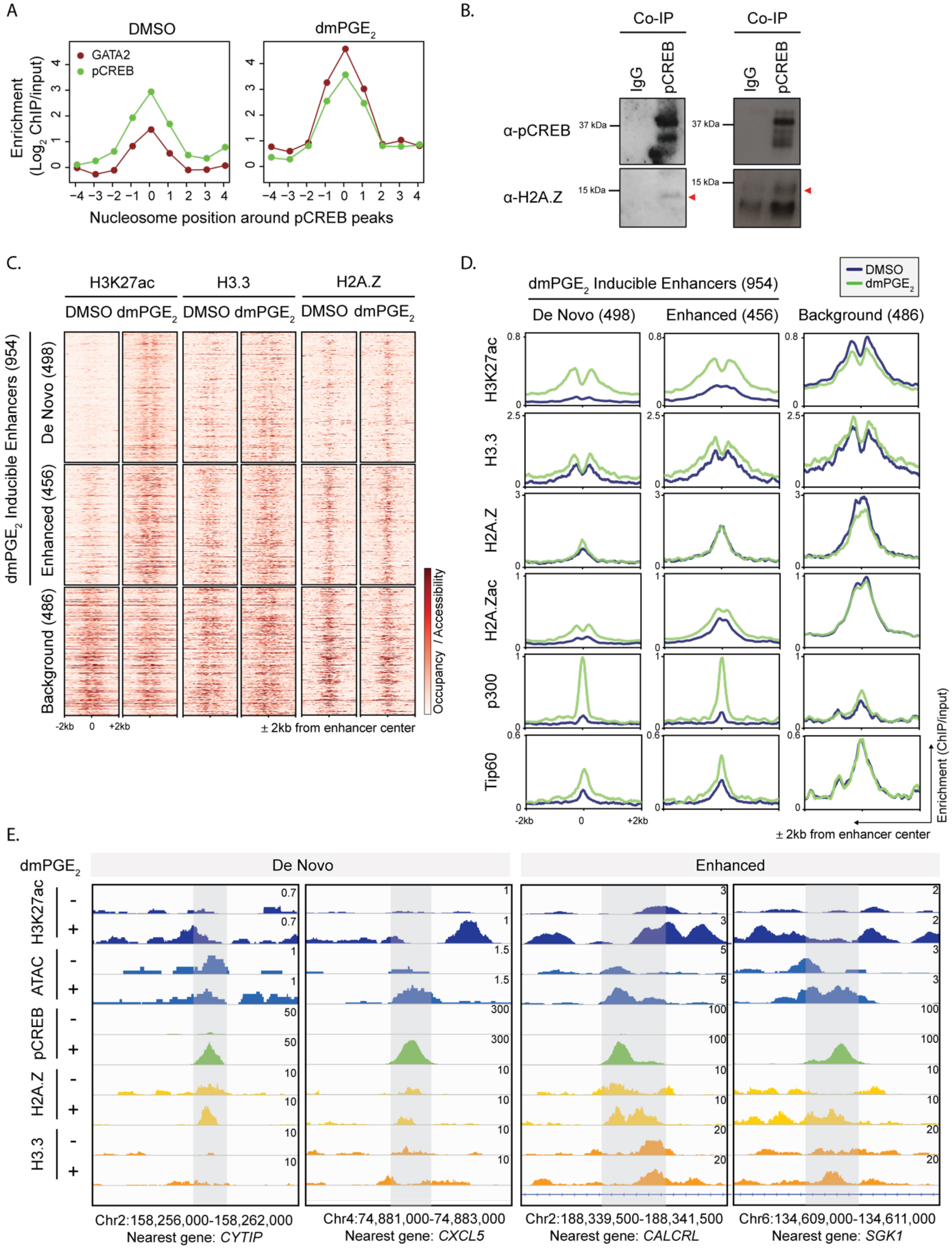

**Supplemental Figure 7. Enhancer nucleosomes contain histone variants.** (A) Enrichment of transcription factors at nucleosome positions surrounding pCREB peaks within enhancers before and after dmPGE$_2$ stimulation. Position 0 indicates the nucleosome overlapping with pCREB peak centers. (B) Complex immunoprecipitation (Co-IP) showing that pCREB associated with H2A.Z in U937 myeloid leukemia cells (n = 3 biologically independent experiments). (C) Heat maps of histone variant enrichment around enhancers before and after dmPGE$_2$ treatment. H3K27ac enriched regions identified using ChIP-Seq are classified as De Novo, Enhanced or Background enhancers according to the change in H3K27ac levels observed following dmPGE$_2$ stimulation (n = 2 biologically independent experiments). (D) Average enrichment profiles of histone variants and HATs before and after dmPGE$_2$ treatment in De Novo, Enhanced or Background enhancers. (E) Enrichment of ATAC accessibility, pCREB binding and histone variant deposition in response to dmPGE$_2$ at 4 representative stimuli-response enhancers. Genomic location of presented window and nearest gene are indicated at the bottom of the panel. For all analysis presented in B and D a randomly sampled comparable number of background enhancers (486) is shown.
